# C9orf72-linked arginine-rich dipeptide repeats aggravate pathological phase separation of G3BP1

**DOI:** 10.1101/2023.03.31.535023

**Authors:** Margot Van Nerom, Junaid Ahmed, Tamas Lazar, Joris Van Lindt, Rita Pancsa, Dominique Maes, Peter Tompa

## Abstract

Ras GTPase-activating protein-binding protein 1 (G3BP1) is the key protein driving the formation of cytoplasmic stress granules (SGs) by liquid-liquid phase separation (LLPS). It is a switch-like protein held in a closed and inactive state by intramolecular electrostatic interactions competitively opened by RNA, activating the protein and initiating its LLPS. Here we show that C9orf72-derived arginine-rich dipeptide repeats PR30 and GR30 (R-DPRs) present in amyotrophic lateral sclerosis (ALS) and frontotemporal dementia (FTD), also bind to G3BP1, switching it to an LLPS-competent open state much more effectively than RNA. Whereas RNA binds G3BP1 with micromolar affinity, and cannot initiate LLPS without crowding agents, R-DPRs exhibit a thousand-fold stronger binding to G3BP1, eliciting rapid LLPS even without crowding. The pathogenic effect of R-DPRs is also underscored by the slow transition of R-DPR-G3BP1 liquid droplets to aggregated, ThS-positive states that can recruit the ALS-linked protein hnRNPA2. Deletion constructs and molecular simulations show that R-DPR binding and LLPS are mediated via binding through the negatively charged intrinsically disordered region 1 (IDR1) of the protein, allosterically regulated by the positively charged IDR3. Bioinformatic analyses point to the strong mechanistic parallels of these effects with the interaction of R-DPRs with nuclear nucleophosmin (NPM1) and also suggest that R-DPRs also interact with many other similar nucleolar and stress-granule proteins, extending the underlying mechanism of R-DPR toxicity in cells.

## Introduction

Amyotrophic lateral sclerosis and frontotemporal dementia (ALS/FTD) are fatal neurodegenerative diseases with a complex genetic background [1, 2]. The major genetic lesion observed in inherited (familial) ALS/FTD is the expansion of a hexanucleotide repeat region in the intronic region of gene C9orf72 [1, 3]. The expanded repeat region in such C9-ALS/FTD cases undergoes repeat-associated non-AUG (RAN) translation, giving rise to five different dipeptide repeats (DPRs), which are intrinsically disordered and non-natural protein products linked with disease etiology [4, 5]. There is a developing consensus that the two arginine-rich DPRs (R-DPRs), poly-PR and poly-GR, are highly toxic, that poly-GA is moderately toxic, whereas poly-GP and poly-PA are not toxic to cells [6-8]. Poly-GR and poly-PR impair key cellular processes [9], such as axon development and branching [10], axonal transport [11], nucleocytoplasmic transport [6], protein translation [12, 13], mRNA splicing [14], proteasomal function [15] and the response to endoplasmic reticulum (ER) stress [16]. Whereas these diverse effects may suggest different underlying molecular mechanisms and targeted cellular components, recent results converge on the idea that the primary, possibly unifying mechanism of R-DPR toxicity is the impairment of the functioning of membraneless organelles (MLOs, also termed ribonucleoprotein granules or bimolecular condensates) in affected cells [17].

Besides membrane-bound organelles, such as the nucleus and mitochondria, cellular processes are also organized via dozens of membraneless organelles, such as the nucleoli in the nucleus and stress granules (SGs) in the cytoplasm [18-20]. The timely formation and dispersion of these MLOs is essential for cellular homeostasis and is thought to proceed by spontaneous demixing from a homogeneous solution of macromolecules by liquid-liquid phase separation (LLPS) under appropriate conditions. The resulting condensates are highly dynamic, liquid-like and they carry out diverse emergent functions, such as speeding up biochemical reactions, providing buffering, filtering or exerting force, among others [21-23].

Due to such essential functions, disturbances in the underlying processes are increasingly linked with a broad range of diseases, such as cancer, inflammation, virus infection, and neurodegeneration [9, 19, 24]. Such diseases, also termed “condensatopathies”, represent a possible novel modality of disease etiology and targeting [24, 25]. Along these lines, there is an almost general acceptance that R-DPRs exert their deleterious effects in C9-ALS/FTD through impairing LLPS of nucleoli in the nucleus [26, 27] and SGs in the cytoplasm [12, 26, 28]. Their effects on nucleoli derive from perturbing the LLPS of its primary driver, nucleophosmin (NPM1) [26, 27, 29, 30]. Although evidence is also compelling for the basic perturbation of SGs by R-DPRs, the underlying molecular mechanism(s) of their profound effect are not known.

R-DPRs overexpressed in cells promote SG formation even in the absence of stress, and the resulting SGs are much less dynamic than the ones that form upon more physiological conditions of stress [26, 28]. R-DPRs bind many SG-linked RNA-binding proteins (RBPs) implicated in C9-ALS/FTD, such as G3BP1 and G3BP2 (Ras GTPase-activating protein-binding protein 1/2), Ataxin1, hnRNPA1 (heterogeneous nuclear ribonucleoprotein A1), FUS (fused in sarcoma), TDP-43 (TAR DNA-binding protein 43), and TIA1 (cytotoxic granule-associated RNA-binding protein) [26, 31], of which G3BP1/2 are essential for SG formation [26, 28, 32, 33]. Several of these RBPs also modulate R-DPR-mediated toxicity in cells [26, 27].

G3BP1 is an essential protein of 466 residues, which has two folded domains (NTF2L and RRM), connected and flanked by long intrinsically disordered regions (IDRs) of opposite net charge. Electrostatic interactions of the negatively (IDR1) and positively (IDR3) charged IDRs keep G3BP1 in a closed, non-phase separating state, while binding of negative RNA to IDR3 and RRM opens up the structure and enables extensive homotypic (G3BP1-G3BP1) and heterotypic (G3BP1-protein and G3BP1-RNA) interactions, promoting SG formation by LLPS upon cellular stress [34, 35]. Under disease conditions, SG formation tends to become irreversible, turning liquid SGs into less dynamic inclusions, a hallmark of ALS/FTD [1, 36, 37]. Here we show that R-DPRs bind very tightly to IDR1 of G3BP1, opening its structure and aggravating G3BP1-driven LLPS, probably providing the basic mechanism of the disruption of SG formation in disease. Their effect is much stronger than that of RNA, and droplets formed by R-DPR driven LLPS undergo a slow transition to aggregated, ThS-positive states. These mechanisms show very strong mechanistic parallels with the effect of R-DPRs on NPM1 in nucleoli [26, 27], which may point to a general R-DPR toxicity mechanism that arises from impairing physiological LLPS driven by many switch-like proteins showcased here by G3BP1 and NPM1.

## Results

### LLPS of G3BP1 is aggravated by Arg-rich C9orf72 dipeptide repeats

G3BP1 acts as a molecular switch that senses the release of mRNA from polysomes stalling upon cellular stress [38]. Upon RNA binding to its positively charged IDR3/RRM region, the structure of G3BP1 opens up and engages in a multitude of protein-protein and protein-RNA interactions, priming the formation of SGs [34, 35]. Aberrant formation of SGs, however, is also a key mechanism of the pathological effect of R-DPRs, of which G3BP1 is a primary target [26, 28]. It occurred to us that the high positive charge of R-DPRs may enable their binding to the negatively charged IDR1 and initiate a switch-like activation mechanism conducive of SG formation, like that elicited by RNA.

To this end, we tested the LLPS of recombinant G3BP1 in the absence and presence of R-DPRs PR30 and GR30, either with or without 1% polyethylene glycol (PEG-10,000), a crowder required for its RNA-driven LLPS [34]. As shown by turbidity measurements carried out with crowder 1% PEG, polyU elicits moderate LLPS of G3BP1, whereas both PR30 and GR30 induce the LLPS of G3BP1 with a signal 20-fold increased (Fig. 1A, Suppl. Fig. S1, S2). These differences are supported by dynamic light scattering (DLS) experiments on the size evolution of LLPS droplets (Fig. 1B). Here, polyU induces the rapid formation of very small droplets (of about 50 nm in diameter) that do not change for about 10 min, in accord with literature suggesting the appearance of small clusters in RNA-G3BP1 phase separation [34]. In contrast, both PR30 and GR30 immediately nucleate large droplets of about 500 nm in diameter, which then evolve toward droplets about 2 µm in size. Striking difference between RNA-and R-DPR-induced LLPS is further highlighted by fluorescence microscopy (Fig. 1C), in which both R-DPRs promote LLPS of G3BP1 very effectively (apparently much more than polyU RNA), while the non-toxic ALS-related DPR, PA30, has no effect. The difference in favor of R-DPRs is even more clear in the absence of PEG, when R-DPRs promote the formation of G3BP1 droplets, whereas polyU RNA has no discernible effect (Fig. 1C). The effective LLPS is apparently promoted by the direct physical interaction of R-DPRs with G3BP1, as fluorescent images show their colocalization in the droplets that form.

**Figure 1:**
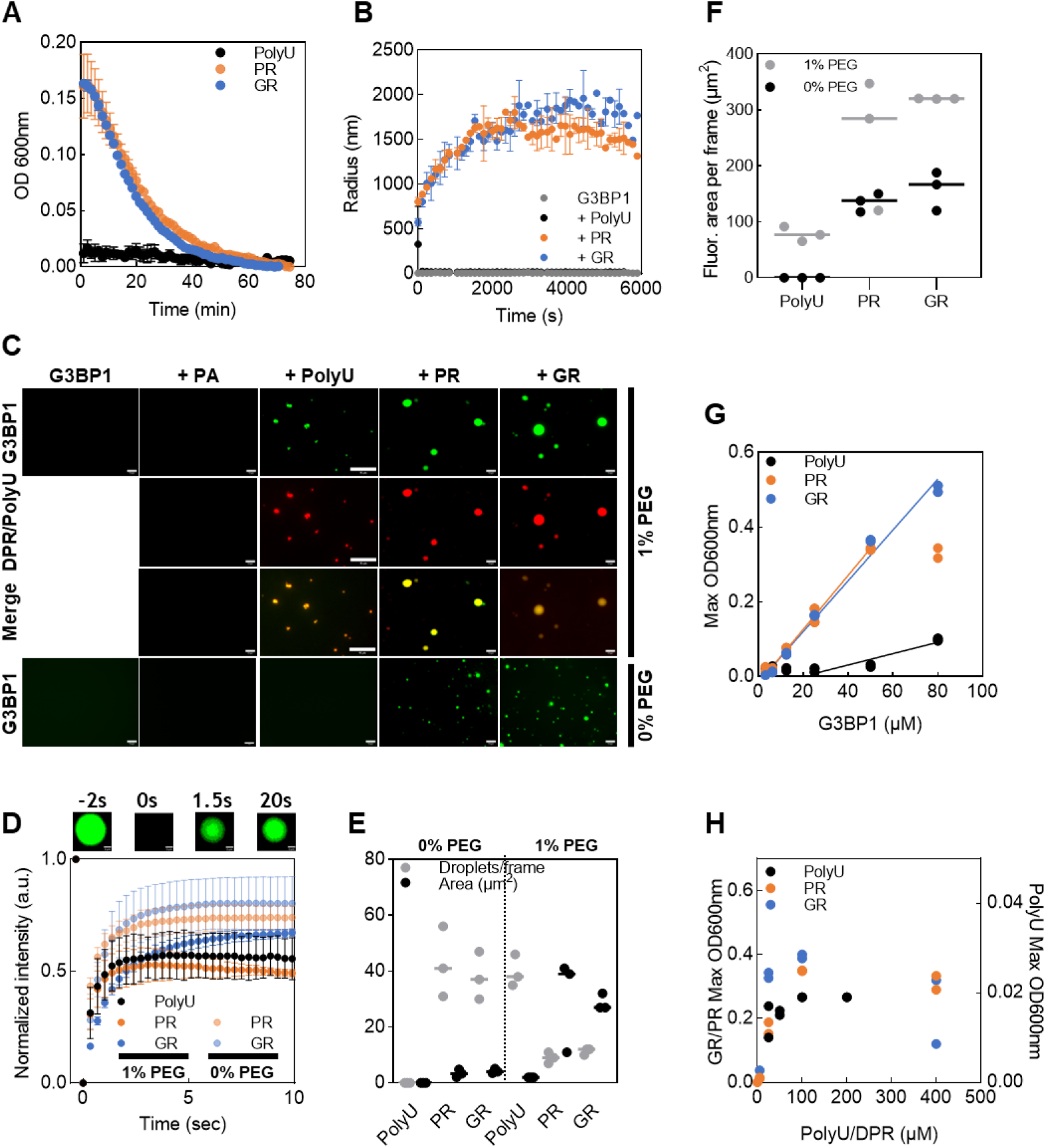
LLPS of G3BP1 is aggravated by Arg-rich C9orf72 R-DPRs. (A) Comparison of phase separation of G3BP1 (25µM) in the presence of polyU (50µM), PR30 (50µM) or GR30 (50µM) followed by turbidity (OD600). (B) The trajectory of the change of droplet size by dynamic light scattering (DLS) under conditions as on panel A. (C) Microscopic images of the LLPS of G3BP1 in the presence (upper panels) or absence (lower panels) of 1% PEG crowding agent. To Alexa-488-labeled G3BP1 at 60 µM, nothing (//), PA30 (PA, 50 µM), polyU (50 µM), PR30 (PR, 50 µM), or GR30 (GR, 50 µM), was added (all partners labeled with Cy5), and microscopic images were recorded after 5 min incubation on ice. The scale bar on all images represents 10 µm. (D) FRAP recovery curves of fluorescence measurements of Alexa-488-G3BP1 droplets (as on panel C), in the presence of polyU, PR30 and GR30, with or without 1% PEG added (shown at t = -2s, 0s, 1.5s and 20s for PR30, 1% PEG, cf. Suppl. Fig. S3 and Suppl. Table S1) (E) G3BP1 droplet statistics of fluorescent images (as on panel Cs): mean and values of 3 repeats of the number of droplets within 3 observation areas, and mean and values of 3 repeats of the surface area of the droplets, shown with or without 1% PEG added. (F) Total fluorescent surface area per frame (number of droplets x their average surface) on panel C. (G) Determination of Csat of LLPS of G3BP1, titrating polyU (50 µM), PR30 (50 µM) and GR30 (50 µM) with G3BP1 and plotting the maximum value of the OD600 turbidity values (Suppl. Fig. S1). Extrapolation of the linear rising phase yields Csat values of 25 µM (polyU) and 2 µM (both PR30 and GR30). (H) Concentration effects of polyU, PR30 and GR30 on the LLPS (maximum OD600 in turbidity measurements) of G3BP1 (25 µM) (Suppl. Fig S2).

The mechanism of condensate formation by LLPS, i.e. the liquid nature of droplets formed is evidenced by fluorescence recovery after photobleaching (FRAP) experiments, in which the fluorescence of Alexa-488-labeled G3BP1 recovers within a few seconds with all the partners used (polyU, PR30 and GR30; see Fig. 1D, Suppl. Fig. S3). Interestingly, some differences appear: (i) in the presence of 1% PEG, the amplitude of recovery in all three systems is only about 50%, whereas in the absence of PEG, DPR droplets recover to a higher extent, about 70%, and (ii) GR30-induced droplets appear less dynamic than PR30-induced droplets, recovering somewhat more slowly, both in the absence and presence of PEG, which may indicate differences in their viscosity (Suppl. Table S1).

When quantifying the microscopic images (Fig. 1C), G3BP1 LLPS dynamics are very different in the presence of R-DPRs as compared to LLPS induced by polyU RNA (Fig. 1E). In the presence of 1% PEG, polyU nucleates about 3 times more droplets than R-DPRs, but these droplets tend to be about 20 times smaller in diameter and/or mature much slower than those forming in the presence of R-DPRs (droplets are about 5 µm radius in the case of R-DPRs, but only 0.2 µm in the case of polyU, after about 5 min incubation time). The difference is even more striking in the absence of PEG, where no LLPS occurs with RNA, even at high protein and RNA concentrations. When determining the total fluorescence signal of R-DPR-induced condensates (size times their number), it is at least 4 times higher than that of G3BP1 in the presence of polyU (Fig. 1F), in agreement with turbidity (OD600) measurements, which suggest that R-DPR-driven condensation is about 4 times more effective than RNA-driven LLPS (in the presence of crowding, Suppl. Fig. S1, S2).

The dramatic difference between RNA-induced and potentially pathological R-DPR-driven LLPS becomes even more apparent when we compare the saturation concentrations (C_sat_) of G3BP1 required to undergo LLPS under the different conditions. By plotting the maximum of OD600 turbidity values (Suppl. Figs. S1 and S2) upon titrating polyU, PR30 and GR30 with G3BP1 (Fig. 1G), there appears to be a remarkable, and pathophysiologically highly significant reduction in C_sat_ from 25 µM (polyU) to about 2 µM (both PR30 and GR30).

These observations suggest dramatic differences between the LLPS of G3BP1 promoted by RNA and R-DPRs. To explore these further, we have also compared the LLPS behavior of G3BP1 upon titrating it with its partners. In particular, we determined if their binding mode results in a reentrant effect, when LLPS is inhibited at the molar excess of either RNA or R-DPRs, as often observed for the LLPS of ribonucleoproteins [39, 40]. Of probable relevance to the molecular mechanisms of LLPS, here we see (Fig. 1H) that polyU and PR30 have no reentrant effect, while GR30 has some minor reentrant tendency, i.e., the magnitude of LLPS slightly decreases above its 5x molar excess to G3BP1.

### A switch-like activation of G3BP1 entails aggravated LLPS

It is suggested that the domain structure and 3D spatial organization of G3BP1 are intimately linked with its RNA-promoted LLPS [34, 35]. G3BP1 contains two folded (NTFL2 and RRM) and three intrinsically disordered (IDR1, IDR2, IDR3) domains (Fig. 2A). Of these, IDR1 has a high negative net charge whereas IDR3 is highly positively charged (Fig. 2A). The molecule is held in a closed conformation by electrostatic interactions between IDR1 and IDR3 (Fig. 2B), and RNA can open the structure and induce LLPS by binding to IDR3 and RRM [34, 35]. We hypothesized that a mechanistically similar mechanism can be initiated by R-DPRs binding to IDR1 (Fig. 2B).

**Figure 2:**
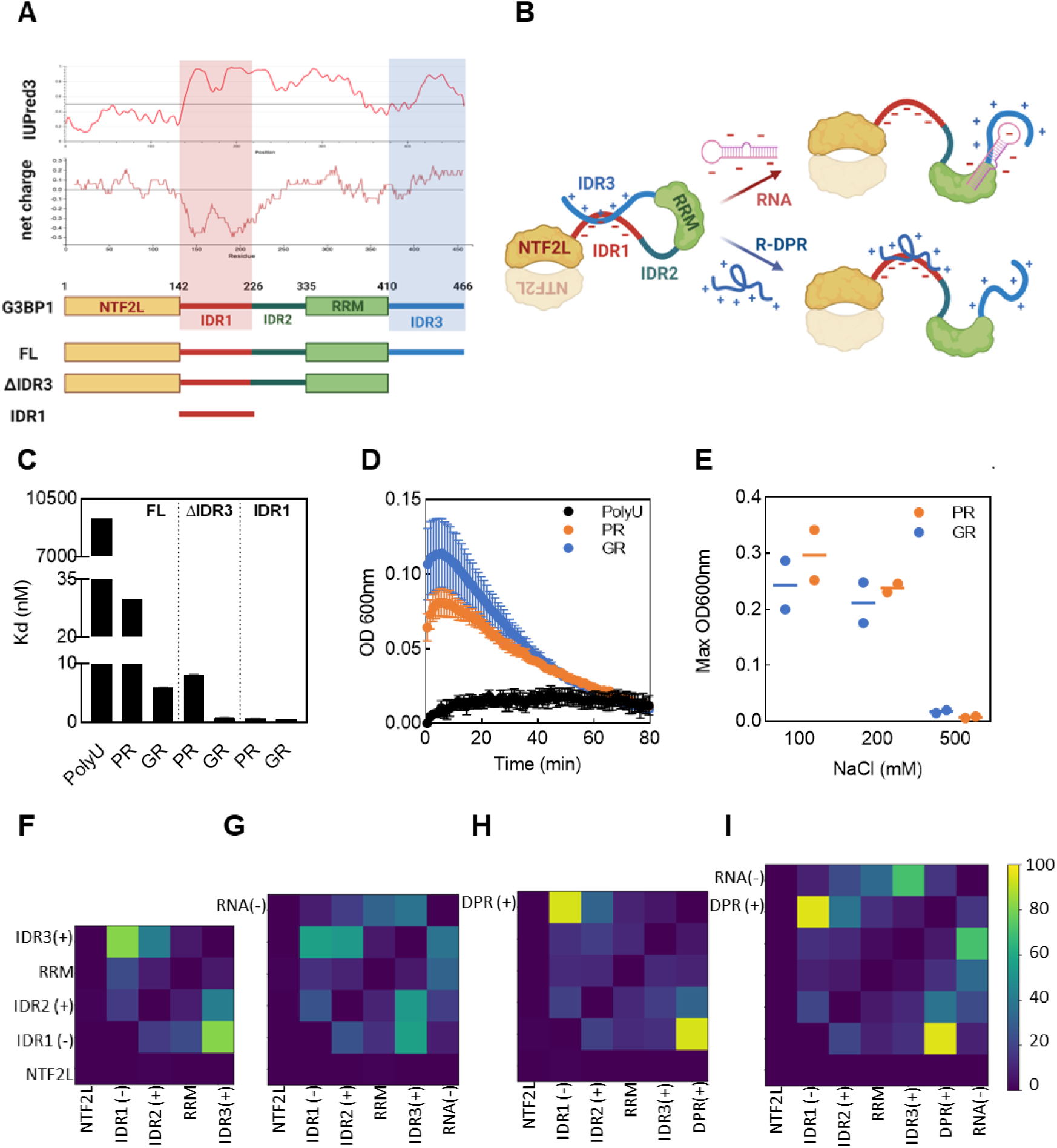
Switch-like activation mechanism of G3BP1 by R-DPRs. (A) Scheme of the domain organization of G3BP1, overlaid with structural disorder predicted by the IUPred algorithm and net charge calculated within a sliding window of 20 residues. Folded (NTFL2 and RRM) and intrinsically disordered (IDR1, IDR2, IDR3) domains are marked, with (negatively charged) IDR1 and (positively charged) IDR3 highlighted. Constructs used in this study are shown. (B) Structural scheme of G3BP1, kept in a closed conformation by electrostatic interactions between IDR1 and IDR3 [34, 35]. RNA binding to ID3/RRM opens the structure; a similar activation mechanism is suggested to occur by R-DPR binding to IDR1. Dimerization is indicated only by a second copy of the NTF2L domain. (Figure 1A and 1B created by BioRender). (C) Binding of polyU, GR30 and PR30 to G3BP1, G3BP1-ΔIDR3 (ΔIDR3) and G3BP1-IDR1 (IDR1): Kd values were measured by BLI (Suppl. Fig. S4, Table S2). (D) LLPS of G3BP1-IDR1(25µM) in the presence of polyU (25µM), PR30 (25µM) or GR30 (25µM) by OD600. (F-I) Region-region contact maps of G3BP1 domains and its partners from ultracoarse-grained single-chain Monte Carlo simulations of single molecules, carried out for a monomeric G3BP1 (F), a G3BP1 in complex with an RNA (G) or a DPR (H), or both an RNA and a DPR molecule (I).

Colocalization experiments (Fig. 1C) in fact suggest a direct physical interaction between G3BP1 and R-DPRs in driving LLPS. Therefore, we first addressed if R-DPRs engage in direct interaction with G3BP1, by applying bio-layer interferometry (BLI; Suppl. Fig. S4) and microscale thermophoresis (MST; Suppl. Fig. S5) to determine the Kd of binding of both R-DPRs and RNA to G3BP1. As shown (Fig. 2C), the binding of both GR30 (Kd = 5.9 ± 0.8 nM with BLI, 19 ± 1.0 nM with MST) and PR30 (Kd = 29.8 ± 0.5 nM with BLI) is very tight, much stronger than with polyU RNA (Kd = 9.3 ± 0.6 µM with BLI, 10.48 ± 1.00 µM with MST; all Kds measured or collected from the literature are presented in Suppl. Table S2).

Next, we asked if binding is driven by the negatively charged IDR1, in a manner similar to RNA binding, which is mediated by the positively charged IDR3 and the adjacent RRM [35]. To this end, we generated two constructs: G3BP1-IDR1 and G3BP1-ΔIDR3 (Fig. 2A) and measured their R-DPR binding strength. The sub-nM binding of G3BP1-IDR1 (Fig. 2C, Table S2) provides direct evidence that this region is the primary driver of the R-DPR - G3BP1 interaction. This is further supported by the removal of the negatively charged IDR3, which makes R-DPR binding significantly more tight than that of the full-length (FL) protein (Kd G3BP1-ΔIDR3, PR30 = 8.1 ± 0.5 nM, GR30 = 0.8 ± 0.4 nM, cf. Fig. 2A and Suppl. Table S2). This much stronger binding provides strong support for the activation model, in which competition between IDR3 and R-DPRs regulates the opening of G3BP1 structure and LLPS of the protein (Fig. 2B), which also manifests itself in the effective LLPS of IDR1 (Fig. 2D) and G3BP1-ΔIDR3 together with R-DPRs. In support of the predominantly electrostatic nature of R-DPR - G3BP1 interaction, we observe it is very sensitive to salt (Fig. 2E, Suppl. Fig. S6), whereas 1,6-hexanediol, known more to interfere with hydrophobic interactions, has a much smaller effect (Suppl. Fig. S6).

As shown by previous SAXS experiments and Monte Carlo simulations [34, 35], RNA brings the molecule from a compact structure to an extended state compatible with RNA-mediated clustering and crosslinking (Fig. 2B). The binding experiments underscore that a similar mechanism is feasible by the strong electrostatic interaction between R-DPRs and IDR1, however, SAXS experiments to provide direct evidence cannot be carried out, due to the strong phase-separation tendency of G3BP1 with R-DPRs. Therefore, to approach the underlying mechanism, we have carried out coarse-grained Monte Carlo simulations of a G3BP1 monomer (not considering its dimerisation via the NTF2 domain) alone (Fig. 2F) or in the presence of an RNA (Fig. 2G) or DPR (Fig. 2H) molecule (for parametrization of the coarse-grained model, cf. Suppl. Table S3). The simulations underscore that: (i) in the absence of a binding partner, the predominant interaction is between IDR1 and IDR3, with secondary interactions between IDR2 and IDR3, and IDR1 and RRM, in accord with the closed conformation suggested (Fig. 2B); (ii) in the presence of RNA, IDR3 - RNA interactions lessen the IDR1-IDR3 interaction; (iii) DPR has a much stronger effect: the IDR1 - DPR interaction becomes dominant, completely abrogating the IDR1 - IDR3 interaction. Most interestingly, when both DPR and RNA are present (Fig. 2I), the IDR1 - DPR interaction is still dominant, but the IDR3 - RNA interaction becomes much stronger than with RNA alone (cf. Fig. 2I vs. Fig. 2G), suggesting an allosteric coupling between IDR1 and IDR3, resulting in positive cooperativity between DPR and RNA binding. This cooperativity is confirmed by direct binding experiments, in which the binding of GR30 to G3BP1 (Kd = 5.9 ± 0.8 nM) becomes even stronger (Kd = 0.02 nM) in the presence of RNA (cf. Suppl. Fig. S4h, Suppl. Table S2).

### Mechanism of G3BP1 LLPS induced by R-DPRs

These results underscore that direct R-DPR - IDR1 binding is responsible for the opening of G3BP1 structure and initiating LLPS of the protein. Previously, it was shown that RNA binds IDR3/RRM, and promotes the formation of local G3BP1-RNA clusters, which are crosslinked by long RNA bridges to promote LLPS [34]. It is, therefore, a reasonable hypothesis that R-DPR binding at IDR1 and RNA binding at IDR3 work by a similar mechanism, leaving the question open if the nature of long-range contacts conducive to LLPS are also similar. We next addressed these issues.

First, we have checked if mutant G3BP1, lacking the IDR3 region, can undergo LLPS. Intriguingly, at a lower concentration than in the previous experiments (12.5 µM), LLPS of FL G3BP1 is not apparent with microscopy (as R-DPRs are at 10 µM, as compared to 60 µM in Fig. 1G), while G3BP1-ΔIDR3 undergoes discernible phase separation (Fig. 3A). This result suggests that the positively charged IDR3 is inhibitory to R-DPR-induced LLPS, i.e., it does not provide crosslinks within the condensates, as later also shown by coarse-grained molecular simulations. The situation may be somewhat different in the presence of RNA: when both R-DPR and RNA are present, their effects are more than additive, showing a strong cooperativity (Fig. 3B). This is, in a way, the consequence of the mutually reinforcing effect of RNA and R-DPR binding on each other, as indicated by the stronger binding of GR30 in the presence of RNA (Fig. 2J).

**Figure 3:**
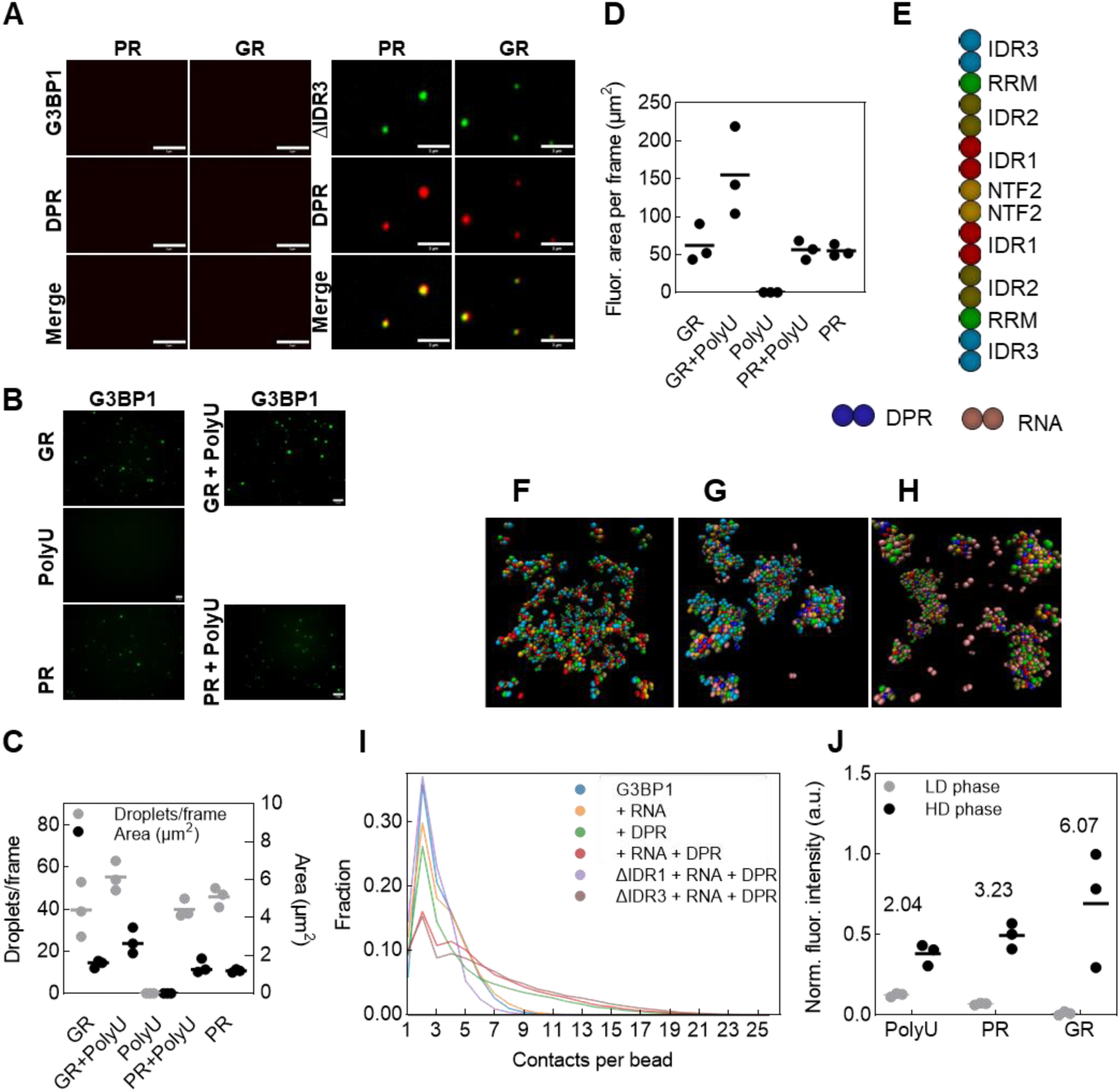
Mechanism of R-DPR - promoted LLPS. (A) Effect of deleting IDR3 on the LLPS of G3BP1: microscopic images of G3BP1 (12.5 µM, left panel) or G3BP1-ΔIDR3 (12.5 µM, right panel) droplets in the presence of PR30 (10 µM) or GR30 (10 µM). (B) Cooperative effect of DPRs and RNA on LLPS: microscopic images of G3BP1 (50 µM) droplets in the presence of polyU (25 µM), GR30 (25µM) or PR30 (25 µM) alone, or polyU + GR30 or polyU + PR30. (C) Statistics of fluorescent images (as on panel B): mean and values of 3 repeats of the number of droplets within 3 observation areas, and mean and values of 3 repeats of the cross-section surface area of the droplets, and (D) total fluorescent surface area per frame (number of droplets x their average surface). (E) Coarse-grained architecture of G3BP1 dimer (held tightly together by two NTF2 domains), DPR and RNA shown with beads matching panels F-H. Snapshots of ultracoarse-grained multi-chain Monte Carlo simulations of 200 G3BP1 dimers (F), 200 G3BP1 dimers, 200 DPR and 200 RNA molecules (G), and 200 G3BP1-ΔIDR3 dimers, 200 DPR and 200 RNA molecules (H). (I) For all states of the on-lattice Monte Carlo simulation trajectories, the number of contacting other beads for every G3BP1 bead was counted; the normalized distribution of beads with various contact numbers is shown. (J) Enrichment of G3BP1 in droplets induced by polyU, PR30 or GR30 (as in Fig. 1C): fluorescence ntensity of droplets (high-density phase) vs. area within droplets (low-density phase).

Such cooperativity supports that R-DPRs and RNA might operate by the same mechanism (Fig. 2B). Upon observing microscopic images of condensates that form in the presence of RNA or a DPR alone, or in combination (Fig. 3B), this picture is primarily substantiated in the case of GR30. When counting the number of droplets or their mean cross-section area (Fig. 3C), or the total area covered by droplets (Fig. 3D), their numbers are additive in the case of PR30, but more than additive with GR30. Also arguing for a mechanism shared by R-DPR and polyU, cooperativity is not seen when both partners are present at saturating concentrations (Suppl. Fig. S7), which would not be the case if they induced additive mechanisms.

To explore further the possible mechanism of LLPS, we performed ultracoarse-grained Monte Carlo simulations on a large number of molecules, with polymers coarse-grained by associating a single bead with the globular domains of G3BP1 and two beads with its disordered regions, as well as DPRs and RNA molecules (Fig. 3E). We have carried out simulations of 200 FL G3BP1 molecules (enforcing dimerization via the NTF2 domains, Fig. 3F), with either 200 RNA or 200 DPR molecules alone (Suppl. Fig. S8), or together (Fig. 3G). The same number of mutants G3BP1-ΔIDR1 (Suppl. Fig. S9) or G3BP1-ΔIDR3 (Fig. 3H) was also simulated in the presence of RNA and DPR molecules. The ensuing trajectories were analyzed for the distribution of contact density, which is a good metric of the condensed state (Fig. 3I). The trends of different combinations suggest that: (i) G3BP1 alone, or together with RNA, has little tendency to undergo LLPS, (ii) R-DPRs promote G3BP1 LLPS much stronger than RNA, (iii) R-DPRs and RNA cooperate, i.e. have a stronger effect together than either alone, (iv) IDR1 is critical for the observed LLPS, as G3BP1-ΔIDR1 does not phase separate even in the presence of DPRs and RNA, (v) G3BP1-ΔIDR3, however, readily phase separates in the presence of DPRs and RNA, with RNA not being involved in condensates, underscoring the secondary importance of IDR3 in R-DPR-driven LLPS.

By analysing detailed domain-domain or domain-partner contact maps of the multi-chain trajectories (Suppl. Fig. S10), the picture gains some more interesting details: (i) the dominant interaction driving LLPS in the absence or presence of RNA is between DPR and IDR1, (ii) when DPR is present, there is always - irrespective of the presence of RNA - there is significant IDR1 - IDR1 interaction, (iii) when LLPS is driven by DPRs, there is discernible interaction – or rather proximity – between IDR1 domains, and also bound DPRs, of adjacent G3BP1 molecules, suggesting a tight packing of G3BP1 molecules in the condensate, (iv) even in the presence of DPR and RNA together, these interactions prevail, and there is also a significant interaction between RNA and DPR molecules.

In all, these analyses suggest that G3BP1 fails to drive LLPS as long as IDR1 is predominantly in contact with IDR3, but if DPRs outcompete IDR3 for IDR1, or if IDR3 is deleted from the construct, G3BP1 will effectively phase separate. RNA alone apparently only masks a certain subpopulation of IDR3 from IDR1, resulting in only partial competition, which is apparently insufficient for G3BP1 condensation. For this system, mostly tight stacking between IDR1 and DPR layers, possibly also including RNA but not that much IDR3, drives phase separation. This is in line with the observed effective LLPS of G3BP1-IDR1 with R-DPRs (Fig. 2D).

The previously suggested RNA-driven crosslinking model via the RRM and IDR3 [34] suggests a low, ∼1 mg/ml G3BP1 concentration in condensates, which is scaffolded mostly by long RNA chains. Our results suggest a different scenario under potentially pathological conditions in the presence of R-DPRs, when close stacking of G3BP1 and R-DPR molecules entails a much higher effective protein concentration in the condensates. To check on this prediction of the model, we assessed the enrichment of G3BP1 in condensates in previous microscopic images of G3BP1 LLPS in the presence of polyU, PR30 and GR30 (Fig. 1C): the ratio of intensities in droplet vs. solution is much higher for R-DPRs than for RNA (Fig. 3J), about 2 (polyU), 3 (PR30) and 6 (GR30), which suggests a much tighter packing of proteins in the latter.

### Pathological implications of R-DPR - G3BP1 interactions

Our results that R-DPRs are order(s) of magnitude stronger than RNA in promoting LLPS of G3BP1 and promote a very different molecular organization, appear to draw strong mechanistic parallels with the profound differences in SG formation under physiological and pathological conditions. RNA-driven physiological SGs are reversible and disperse readily upon the cessation of stress [36], whereas SGs forming under pathological conditions tend to persist and turn into increasingly viscous gel-like states, eventually giving rise to aggregates [41], as readily induced by repetitive stress [37]. This observation is thought to be conducive of cellular inclusions forming upon the action of R-DPRs, both in cellular expression systems and in ALS/FTD-linked pathology [4].

To follow up on these potential mechanistic parallels with cell physiology and pathology, we have tested and compared the behavior of G3BP1 - polyU and G3BP1 - R-DPR droplets. A hallmark of C9-ALS/FTD is the appearance of R-DPR-positive inclusions/aggregates, which are often characterized by ordered, amyloid-like states positive for thioflavin T (ThT) fluorescence (substituted by Thioflavin S (ThS) in the presence of RNA), as also demonstrated by SG induction by R-DPR overexpression in cells, for example [28]. To this end, we measured ThS fluorescence for G3BP1, and G3BP1 mixed with polyU, PR30 or GR30, incubated over almost five days (Fig. 4A). The differences are striking: R-DPRs cause a transition to a ThS positive state over time, much more than polyU, or G3BP1 alone. We have also followed the possible differences by fluorescence microscopy (Fig. 4B). In the absence of partners, or the presence of polyU, no aggregates are formed over 4 days of incubation.

**Figure 4:**
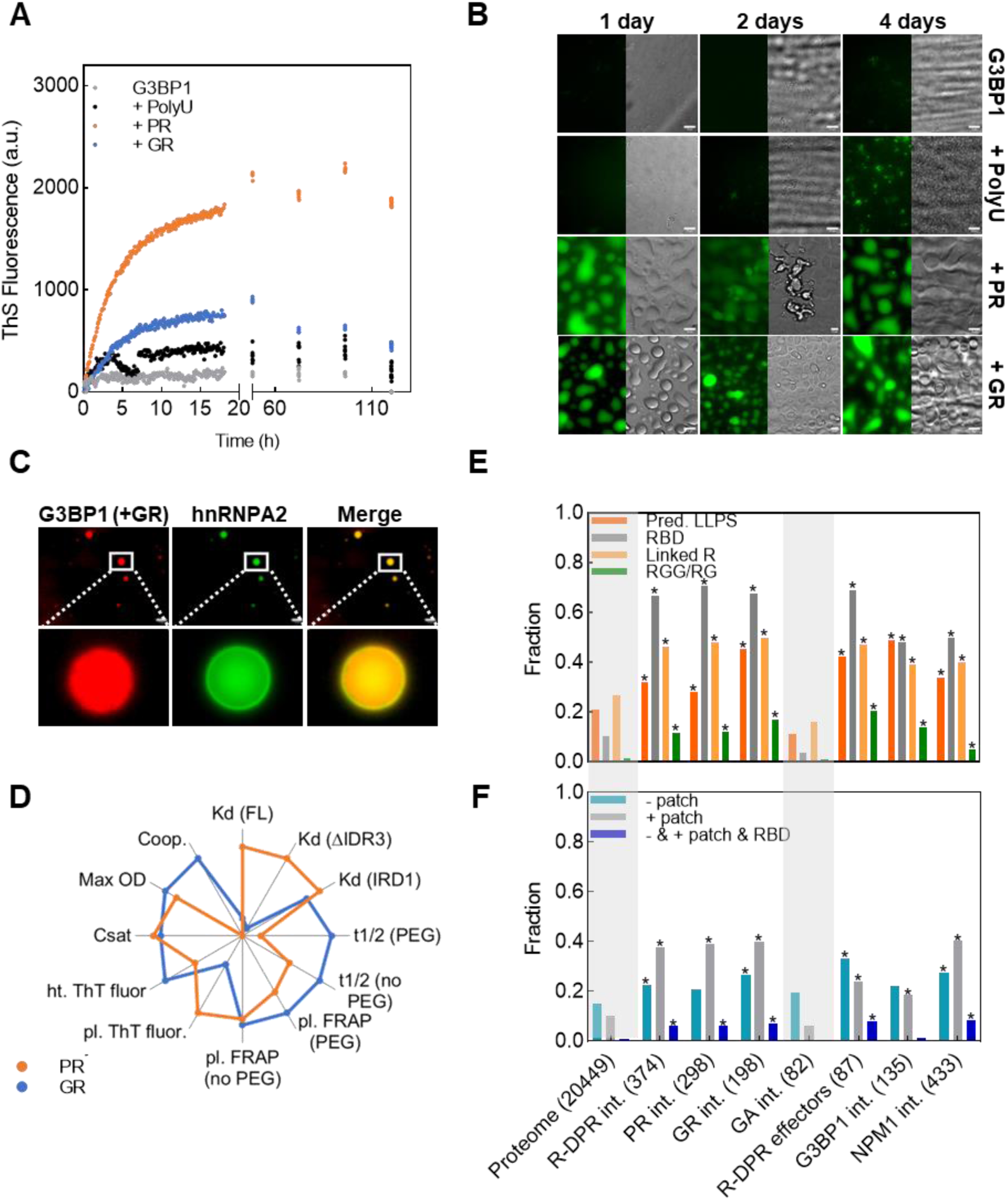
R-DPR promoted G3BP1 LLPS features disease aspects of C9-ALS/FTD. (A) Change in the material state of G3BP1 condensates over time: ThS fluorescence of G3BP1 (60µM) alone, or in the presence of polyU (50 µM), PR30 (50 µm) or GR30 (50 µM). (B) Microscopic images of G3BP1 (60µM) droplets in the presence of polyU (50µM), PR30 (50µM) or GR30 (50µM), after incubation of 1, 2 and 4 days in presence of ThS. (C) G3BP1-R-DPR condensates recruit hnRNPA2. Microscopic images of G3BP1 (30µM, red) in presence of GR (30µM) and hnRNPA2 (green). (D) Star diagram outlining the differences in PR30-and GR30-promoted G3BP1 condensation: Kd values of PR/GR with G3BP1, G3BP1-ΔIDR3 or G3BP1-IDR1 (Fig. 2C); plateau (pl.) and halftime (t1/2) of FRAP measurements in presence or absence of PEG (Table S1, Fig. 1D) and of ThT measurements (Fig. 4A); MaxOD measurements and derived C_sat_ (Fig. 1G); Cooperativity for GR, no cooperativity for PR. (Fig. 3D). In all comparisons, values have been normalized to the highest value being 1. (E) Fraction of proteins in the whole human proteome and among the interaction partners of R-DPRs (polyPR and polyGR also shown separately, also “effectors”, i.e., partners which affect R-DPRs toxicity [26]), polyGA, G3BP1 and NPM1, possessing various features: LLPS propensity by DeePhase (threshold 0.75), or containing an RNA-binding domain (RBD), two linked R motifs, or an RGG repeat region. (F) Fraction of the same proteins shown to contain a negative patch (≥ –10 net charge w/in 30 amino acids, -patch), a positive patch (≥ +10 net charge w/in a window of 30 amino acids, +patch), or three features combined: a negative patch, a positive patch and an RBD (-& + & RBD). In panels E and F, bars of significant difference from the respective proteome value by chi2 tests are marked by a star. Bonferroni correction was applied on the significance levels (they were reduced tenfold): p<0.005 is marked *. For exact p-values and significance levels, see Suppl. Table S6.

Formation of physiological SGs is compromised in C-9ALS/FTD, also by the pathological recruitment of mislocalized, additional ALS/FTD-linked RBPs, like TDP-43, FUS, hnRNPA2, TIA1 and others [41]. To appreciate the disease-linked features of G3BP1 phase separation induced by R-DPRs, we next asked if hnRNPA2 is recruited into these condensates. We added DyLight 488-hnRNPA2 to the G3BP1 condensates forming in the presence of GR30. As hnRNPA2 also has a strong tendency to phase separate, we kept it under its own C_sat_ [42] and visualized its location by fluorescence microscopy (Fig. 4C). Clearly, it has a strong tendency to partition into R-DPR-G3BP1 condensates, which may be relevant with disease. Whereas distribution of G3BP1 appears homogeneous in the droplet formed, hnRNPA2 is enriched around its periphery, recapitulating earlier observations that collective interactions of proteins and RNA are sufficient for driving the formation of multilayered condensates with distinct material properties [43], which is also a key feature of the core-shell architecture of cellular SGs [44].

An additional, little appreciated aspect of the effect of R-DPRs is that the two, poly-PR and poly-GR, are often indiscriminately mentioned together, although they must be structurally very different (Gly being very flexible and Pro favoring rigid, extended conformations) and many observations in fact underscore their differences, probably playing disparate roles in ALS/FTD [4]. To survey if our observations support differences between the two R-DPRs, we have collected and compared all quantitative data on their behavior (Fig. 4D). There are non-negligible differences between, for example in terms of their Kd of G3BP1 binding, t1/2 of FRAP recovery, cooperativity with RNA in driving LLPS, etc…

### Interactomes of R-DPRs underscore their roles in impairing stress granule and nucleolar function

To get further insight into the intricacies of the pathological functions of R-DPRs, we next carried out a detailed analysis of their cellular interactomes. It is to be emphasized that unlike functional cellular proteins, R-DPRs are highly non-natural pathological products of erroneous translation, with non-evolved interactomes, which thus must reflect strongly on their pathological functions. To gain insight on these, we have compiled a list of interaction partners of poly-PR (298), poly-GR (198), both R-DPRs (374), G3BP1 (135), and NPM1 (433) from high-throughput studies [26, 45, 46] and the BioGRID interaction database (for details, cf. Methods). As controls, we have taken the entire human proteome (20449), the interaction partners of ALS/FTD-linked non-toxic poly-GA (82), and also a subset of R-DPR interactors (87), whose homologs showed a strong effect on R-DPR toxicity in a Drosophila RNAi silencing experiment (enhancers or suppressors, hereafter called R-DPR “effectors”) [26] (all data are in Suppl. Table S4).

First, upon comparing the cellular binding partners of poly-GR and poly-PR, although their interactomes overlap to a great extent (Suppl. Fig. S11a), their unique, non-overlapping, interactors show characteristic differences: poly-GR shows some preferences for SGs: 19/76 (25%) of their unique interactors are involved with SGs (only 19/176 (10.8%) for poly-PR), whereas the same numbers are 5/76 (6.6%) for poly-GR with nucleoli, and 23/176 (13.1%) for poly-PR.

A further, general pathological angle of our observations emanates from the switch-like electrostatic activation of G3BP1, whether promoted by RNA or R-DPRs (Fig. 2B). This activation mechanism shows strong mechanistic parallel with the effect of RNA/R-DPRs on the nucleolar protein nucleophosmin (NMP1), the major nucleolus driver apparently targeted by R-DPRs in ALS/FTD. Of note, NPM1 (i) also has acidic and basic IDR tracts and an RNA-binding region (RBD), which engage in intermolecular electrostatic interactions, (ii) these interactions are broken by RNA under physiological, and R-DPRs under pathological conditions [26, 27, 29, 30]. To address if the robust effect of R-DPRs on G3BP1 and NPM1 may showcase some more general pathological mechanisms, we have taken a broad comparative analysis of the interactomes of R-DPRs, G3BP1 and NPM1 (cf. Suppl. Tables S5 and S6).

First, we asked if the interacting proteins tend to undergo phase separation. By an LLPS driver predictor, DeePhase [47], we found that R-DPR-, G3BP1-and NPM1 interactors are significantly more LLPS prone than the human proteome (Fig. 4E). Importantly, poly-GA interactors are less LLPS prone (chi2 test; p=0.00012) than R-DPR interactors. Next, we asked if the interaction partners have features relevant to binding RNA, or negatively charged sequence patches, as identified for NPM1 partners [26, 27]. Interestingly, the presence of an RBD, linked R motifs (at least two RxR or RxxR motifs located maximum 10 residues apart), or linked RGG/RG motifs are all enriched not only in NPM1 and G3BP1, but also in R-DPR partners (chi2 test, p<0.001 for each). Also, significantly less poly-GA interactors show these features than R-DPR interactors (chi2 test; p<0.005 for each feature). These strong preferences suggest very strong mechanistic parallels between NPM1 and G3BP1 (statistical comparisons of calculated features are in Suppl. Table S6).

The switch-like activation of G3BP1 [34, 35] and NPM1 [26, 27] relies on the presence of intrinsically disordered negative and positive patches that can engage in intramolecular electrostatic interactions enabled by IDR flexibility, and RBD(s) that enable RNA to play an active role in switching the protein (cf. Fig. 2B). To find out if R-DPRs may act on a broader set of such inherently switch-like proteins, which could be intimately linked with their toxicity, we looked into these features next (Fig. 4F). As expected, R-DPR partners are enriched in proteins with negative patches compared to the proteome, but unexpectedly, even more in proteins with positive patches that are not their natural partners (chi2 tests, p<1e-4 and p<1e-5, respectively). Of relevance, poly-GA partners do not show any significant differences from the proteome. Most interestingly, R-DPR interaction partners are highly enriched in proteins that show all three features simultaneously (negative patch, positive patch, and an RBD) (chi2 test, p<1e-5), and this combined feature is higher for effectors, which modulate R-DPR toxicity; this is also characteristic of the interaction partners of NPM1.

In all, these strong preferences convey a very important message: not only G3BP1 and NPM1 share similar switch-like features making them vulnerable to R-DPR misregulation, these features are also shared by several other proteins, 23 of R-DPR interactors, and 7 among R-DPR toxicity effectors (Suppl. Table S7). When looking into the list of these predicted proteins, one cannot miss the very strong association with MLOs targeted in ALS/FTD, associated with nucleoli (17), stress granules (8) or both (5). Their even broader toxicity relying on the noted mechanism may be signalled by targeting similar P-body (7) and nuclear speckle (4) proteins. These overlaps are put in context by the similarity of all LLPS-related and switch-like features not only by R-DPR interactors but, surprisingly, also by NPM1-and G3BP1 interactors (Suppl. Fig. S11b). A great proportion of R-DPR interactors (40.1%,150/374) also interact with either G3BP1 or NPM1, while 31.1% (42/135) of G3BP1 interactors and 28.9% (125/433) of NPM1 interactors are also binding partners of R-DPRs. These numbers are very significantly higher than the ones for poly-GA interactors (Suppl. Fig. S11c), which are all vivid manifestations of that R-DPRs hit specifically and heavily the interactome of G3BP1 and NPM1, i.e., SGs and nucleoli.

## Discussion

DPRs produced by C9orf72 hexanucleotide repeat expansion arise as a result of the most prevalent familial mutation in ALS/FTD [1, 3]. There are multiple lines of evidence that poly-PR and poly-GR - the two R-DPRs – exert their toxicity and pathological effect via perturbing the physiological balance of the formation, dynamics and dissolution of cellular MLOs, nucleoli in the nucleus [26, 27] and SGs in the cytoplasm [12, 26, 28]. Whereas their effect on nucleoli apparently proceeds via perturbing the nucleolar driver NPM1 [26, 27], their target and mechanism in SGs has not yet been unveiled. In terms of the underlying molecular mechanisms, NMP1 is kept in a closed, LLPS-incompetent state by electrostatic attraction between its oppositely charged acidic (A) and basic (B) IDRs, and both RNA and R-DPRs can compete off this electrostatic interaction, thereby opening NPM1 structure and enabling a range of homotypic and heterotypic interactions, promoting physiological and pathological LLPS, respectively. We hypothesized that there might be an analogous mechanism in SG misregulation by R-DPRs, as the primary SG driver G3BP1 (and its paralogue G3BP2) also occupies a central role in the SG interaction network [35], and operates by a similar electrostatic switch-like activation between closed and LLPS-competent open states promoted by RNA [34, 35].

In line with these hypotheses, multiple cellular studies imply that G3BP1 could be the prime cytoplasmic target of R-DPRs. First, G3BP1/2 are among the proteins interacting with PR30 in cellular extracts [26, 28]. Overexpression of PR100 in HeLa cells induces very strong SG formation, much stronger than than that by non-toxic PA100, in a strictly G3BP1/2-dependent manner [6]. In another study [26], G3BP1/2 have also been found to be the prime interaction partners of overexpressed PR50 and GR50 and shown to be among the top genetic enhancers of GR50 toxicity in a Drosophila model. Interestingly, similar features have been found for NPM1, the primary driver of RNA-induced formation of nucleoli [26].

All these points implicate G3BP1 as the primary cytoplasmic target of R-DPRs in C9-ALS/FTD. Here we provide compelling evidence for this notion and dissect the underlying molecular mechanisms of G3BP1 activation by R-DPRs, which leads to aggravated LLPS that explains pathological SG induction in C9-ALS/FTD. We do observe that R-DPRs promote the LLPS of G3BP1 much more intensively than RNA, as they can promote condensation in the absence of a crowder, or about 5 times more intensively in the presence of a crowder, and typically at a G3BP1 C_sat_ concentration an order of magnitude lower than RNA. These findings probably parallel, and mechanistically underscore earlier observations [12, 26, 28] of the aggravated formation of SGs in cellular models of R-DPR intoxication. The pathological implications of these differences are also underlined by the notable tendency of R-DPR - G3BP1 droplets to evolve toward ThS-positive states and recruit hnRNPA2, also involved in ALS/FTD inclusions.

These strong effects on LLPS stem from a very strong binding of R-DPRs to G3BP1: their interaction is about 1000 times tighter than that of RNA, whereas binding data with G3BP1- IDR1 and G3BP1-ΔIDR3 constructs, and molecular simulations point to IDR1 being the primary interacting site of R-DPRs. Salt sensitivity underscores the electrostatic nature of the interaction between positively charged R-DPRs and the negatively charged IDR. Prior data suggest that G3BP1 is held in a compact, LLPS-incompetent state by attractive electrostatic interactions between its two long IDRs, the negatively charged IDR1 and positively charged IDR3 [34, 35]. Under physiological stress, when mRNA is released from stalled polysomes, it binds to IDR3, opens G3BP1 structure and promotes LLPS by extensive protein-protein, protein-RNA and RNA-RNA interactions [34, 35]. Droplets thus formed in vitro, and SGs probably driven by an analogous mechanism in the cell, are highly dynamic, showing rapid recovery by FRAP and rapid dissolution upon the termination of stress.

Stark differences of the mechanism of LLPS with RNA and R-DPRs underscore the distinction between physiological and pathological situations, as the LLPS promoted by RNA [34] has many features different from that we observed for R-DPRs. Whereas a short RNA (A60) cannot make G3BP1 phase separate, R-DPRs of only 30 dipeptide units do, and even with long RNA (e.g., total cell extract [34]) LLPS requires crowding, not needed with R-DPRs. In addition, whereas RNA-based LLPS requires many different protein-protein, protein-RNA (and probably RNA-RNA) interactions, as G3BP1 constructs ΔNTF2L and ΔIDR3 (termed originally ΔRG) do not phase separate, suggesting that besides RNA-RRM interactions, NTF2L-NTF2L, IDR3-IDR3, RNA-IDR3 and RNA-RNA interactions all play a role [34]. In the case of R-DPRs, however, even G3BP1-IDR1 and G3BP1-ΔIDR3 can effectively phase separate with R-DPRs, suggesting a very different mechanism, where the IDR1 - R-DPR interaction is sufficient for driving LLPS. With RNA, condensates form by small RNA-G3BP1 clusters crosslinked by long RNA molecules forming an extended network [34], resulting in condensates with high (64 mg/ml) RNA, but very low G3BP1 (1 mg/ml) concentrations. In contrast, with R-DPRs, molecular simulations suggest rather dense protein condensates, driven by IDR1 - R-DPR - IDR1 stacking, explaining why G3BP1 in these R-DPR-driven condensates reaches an order of magnitude higher local concentrations.

A further interesting and highly intriguing aspect of the general perception of pathological R- DPR effects is the difference between the behaviour of poly-GR and poly-PR we observe. Due to their similar, strong toxicity and generation from the same extended hexanucleotide repeat region (poly-GR from the sense, and poly-PR from the antisense mRNA strand [9]), the two arginine-rich DPRs are often mentioned and treated together (as R-DPRs), and results with any one of them are often generalized [4]. Even in the literature, however, there are significant and probably meaningful differences between their pathological actions. For example, when administered externally, GR20 and PR20 polymers enter (U2OS) cells, migrate to the nucleoli, and have a morphological-toxic effect, which is much more pronounced for PR20, which also has much longer cellular half-life (72h) than GR20 (20-30 min) [7]. Whereas poly-PR is suggested to be more toxic than poly-GR [7, 48], it could be due to its much higher cellular stability [7].

Their cellular locations also show characteristic differences. For example, upon transfecting GFP-GR50 and GFP-PR50 into HeLa cells, GFP-GR50 localizes both in the cytoplasm and the nucleus, whereas GFP-PR50 is found predominantly in the nucleus [26]. When measuring diffusion within nucleoli in HeLa cells transfected with mCherry-GR50 and mCherry-PR50, FRAP recovery of nucleolar GFP-NPM1 drops from 60% to 40% in the case of both R-DPRs. When comparing nucleolar GFP-DPR recovery, however, GFP-PR50 recovers to 60%, but GFP-GR50 only to about 20%, suggesting that NPM1 may not be the only target of GR50 in the nucleolus. When cytoplasmic localization is checked, GFP-GR50 colocalizes with SG markers, but GFP-PR50 is not detectable in SGs. In addition, the expression of DPRs in primary motor neurons shows a much stronger nuclear aggregation with PR50 than with GR50 [48]. A further potentially very interesting difference is that in a Drosophila RNAi screen for genetic modifiers of GFP-GR50 toxicity, NPM1 is a strong suppressor, whereas G3BP1 is a strong enhancer of toxicity, again arguing that the primary target of poly-GR toxicity is not NPM1 but G3BP1 [26].

Probably in line with these observations, we do observe a strong effect of both PR30 and GR30 on G3BP1, but characteristic differences between the two R-DPRs. FL G3BP1, G3BP1- IDR1 and G3BP1-ΔIDR3 constructs are all bound much stronger by GR30 than PR30, and more cooperativity is observed between RNA and GR30 than between RNA and PR30. Furthermore, GR30-G3BP1 condensates appear to be more dense than PR30-G3BP1 condensates, and when analysing their cellular interactomes, poly-GR interactome shows some preference for SGs. These findings underscore the observed differences and the above literature distinctions, which refine the potential pathological action of R-DPRs, suggesting a kind of specialization between the two R-DPRs, poly-PR primarily affecting NPM1 in nucleoli whereas poly-GR having more effect on G3BP1 in SGs.

The strong effects of R-DPRs on G3BP1 provide the mechanistic context for findings that overexpression of GR50 and PR50 promotes spontaneous SG formation, without stress, and these SGs are less dynamic, showing an impaired G3BP1 exchange, being also much less inclined to disassemble than the ones induced by stress [26]. As the major pathological hallmark of C9orf72 R-DPR-related ALS/FTD is the appearance of cellular ribonucleoprotein inclusions enriched in R-DPRs and other pathological proteins [4, 5], and a prominent mechanism of R-DPR effects in C9-ALS/FTD is to compromise SG dynamics, it is not far-fetched to conclude that the observed tendency of G3BP1 droplets to recruit the C9-ALS/FTD-related hnRNPA2 is also mechanistically linked to pathological processes characteristic of ALS/FTD. This fits into the general framework of our study, i.e. that a great proportion of R-DPR interaction partners display a switch-like character, and also interact with G3BP1 and NPM1, thus having the potential to have a profound effect on these MLOs.

As a final note, we should not miss that besides ALS/FTD, RAN translation generates potentially aggregation-prone peptide repeats in at least 10 repeat expansion disorders [49] Poly-GP and poly-PR feature in spinocerebellar ataxia 36 (SCA36) [50], poly-EG and and poly-RE in X-linked dystonia parkinsonism (XDP) [51], and poly-LPAC and poly-QAGR in myotonic dystrophy type 2 (DM2) [52]. In addition, repetitive, cationic peptides are used by a broad range of organisms as venoms, toxins and antimicrobials [53, 54], which may similarly target a broad range of host RBPs involved in biomolecular condensation as their primary modality of toxicity. In accord, the mechanistic insight generated here may not only open novel avenues for targeting ALS/FTD, but it may also help better understand these related biological phenomena at the molecular level.

## Material and methods

### G3BP1 constructs

The DNA-construct encoding N-terminally GST-tagged and C-terminally polyHis-tagged full length G3BP1 within a pGEX-2T vector was provided by Prof. Paul-Taylor (St. Jude Children’s Research Hospital, Memphis, TN, USA) [35].

This plasmid was used as a template to generate deletion of IDR3 (aa. 412E-466Q) by using Q5® Site-Directed Mutagenesis Kit with 5’GAAAACCTGTATTTTCAGG3’ forward and reverse 5’TACATTAAGACGTACCTC 3’ primers. Positive clones were sequenced and amplified by transformation in NEB® 5-alpha Competent E. coli. E. coli Rosetta 2 (DE3) were transformed for expression of protein.

The pGEX-2T carrying the fragment for full length G3BP1 was used as a template to generate a plasmid (pHYRSF53 vector) carrying a fragment encoding SUMO-tagged IDR1 through Gibson assembly. Forward primer 5’AAGAAACGGCTCCCGAAGACTAAGCTTGCGGCCGCAC3’ and reverse primer 5’GACTCCTCCTGCGGCTCCGTGGATCCACCAATCTGTTCTCTGTG 3’ for vector amplification and forward primer 5’GGATCCACGGAGCCGCAGGAGGAG3’ and reverse primer 5’AAGCTTGTCTTCGGGAGCCGTTTC3’ were used in combination with NEBBuilder® HiFi DNA assembly.

### Protein expressions and purification

E. coli Rosetta 2 (DE3) carrying the pHYRSF53 vector with the fragment encoding IDR1 or the pGEX-2T vector with fragment encoding full length G3BP1 or the mutant variant G3BP1-ΔIDR3 were grown in LB medium with carbenicillin and chloramphenicol (G3BP1 and G3BP1-ΔIDR3) or kanamycin (IDR1) at 37°C, while shaking until and OD600nm of ± 0.8 was reached. Temperature was then lowered to 16 oC or 28°C (IDR1) and expression induced with 1 mM isopropylthio-β-galactoside (IPTG) overnight. Pelleted cells were then harvested and resuspended in a lysis buffer (50 mM HEPES, pH 7.5, 250 mM NaCl, 1 mM TCEP) complemented with protease inhibitor and DNase, flash frozen and stored at −80 °C.

After thawing, lysis was performed by sonication (Sonics VCX-70 Vibra cell) for 3 min (10 sec pulse on, 10 sec pulse off) at 70% amplification. Next, the sample was pelleted by centrifugation at 4 oC for 1 hour at 19 000 x g.

The supernatant or cell extract was then filtered through a 0.45 µm pore filter and loaded on a HisTrap HP column (GE Healthcare) with immobilized Ni2+ beads equilibrated with HisBufferA (50 mM HEPES, pH 7.5, 250 mM NaCl, 1 mM TCEP and 50 mM imidazole). PolyHis-tagged protein was captured by the column and eluted with a linear imidazole gradient (50 mM imidazole to 500 mM imidazole over 10 column volumes).

Eluted GST-G3BP1, and GST-G3BP1-ΔIDR3 (deleting E411-Q466) was then loaded on a GST HP column (GE Healthcare) equilibrated with GSTBufferA (50 mM HEPES, pH7.5, 250 mM NaCl and 1 mM TCEP). TEV protease was added manually to the column and cleavage of GST-tag and His-tag was performed on column at 4 oC for 4 hours. The sealed column containing the protein was then combined with an equilibrated HisTrap HP column and untagged protein was eluted with HisBufferA. Fractions were pooled and transferred to G3BP1 Storage Buffer (50 mM HEPES, pH7.5, 200 mM NaCl and 1 mM TCEP) through dialysis overnight. G3BP1 and variants were concentrated to 80µM-200µM, flash frozen, and stored at −80 oC until use.

The eluted SUMO-G3BP1-IDR1 (T143-D226) was transferred to a cleavage buffer (50mM HEPES, pH 7.5, 250mM NaCl, 1mM TCEP) and SUMO-tag was cleaved off for 1hr on ice. Sample was loaded on an equilibrated HisTrap HP column and cleaved IDR1 was collected, concentrated to 700µm, flash frozen and stored at −80 °C.

hnRNPA2 was expressed and purified as in [42].

### Dipeptide repeats

Three different DPRs related to ALS/FTD were used. Toxic arginine-rich [Pro-Arg]30 (PR30) and [Gly-Arg]30 GR30, and non-toxic [Pro-Ala]30 (PA30), were purchased from SynPeptide Co., LTD. All three DPRs were dissolved at 1 mM concentration in water, aliquoted, and stored at −80 oC until use.

### Nucleic acid labeling

Polyuridylic acid (polyU, catalog no, P9528-25MG from Sigma-Aldrich) was labeled by using Cy5-conjugated cytidine (bis)phosphate (Jena Bioscience) and biotinylated cytidine (bis)phosphate (Thermo scientific) using T4 RNA ligase enzyme from Thermo Fisher Scientific.

### Fluorescence labeling of proteins

Peptides and G3BP1 at 10 uM concentrations were labeled by using Nanotemper Protein Labeling Kit RED-NHS 2nd Generation (MO-L011) and Alexa Fluor™ 488 NHS Ester (A20100) by following manufacturer protocol. The labeling sample was dialysed overnight with 50 mM HEPES, pH 7.5, 200 mM NaCl and 1 mM TCEP buffer, aliquoted and stored at −80 °C. hnRNPA2 was labeled with DyLight-488 according to the manufacturer’s protocol.

### LLPS assay, turbidity

To follow liquid-liquid phase separation according to turbidity, the optical density (OD) at 600 nm was measured at 25°C for 2 h in a BioTek SynergyTM Mx plate reader while shaking. Samples were prepared in a total volume of 25 µL in a Cell culture 384 well microplate with clear bottom (Greiner). Each experiment was repeated 2 times.

### LLPS assay, microscopy

To visualize droplets, microscopic images were captured by a Leica DFC7000 GT camera connected to a Leica DMi8 microscope. Samples were prepared by initiating liquid-liquid phase separation of green fluorescently labeled G3BP1 in presence of an excess of non-labelled G3BP1 with red fluorescently labeled DPRs or polyU in an excess of non-labelled DPRS or polyU. After incubation of 5 min on ice, the samples were visualized on a microscopic slide with a 100x oil immersion objective. Fluorescence was visualized by a FITC filter (green) and a Rhodamine filter (red).

Experiments with hnRNPA2 LCD were performed similarly with red fluorescently labeled G3BP1 in an excess of non-labelled G3BP1 and green fluorescently labeled hnRNPA2 and non-labelled GR30

To quantify fluorescent images, they were transferred to 8-bit images and analyzed by ImageJ software: of each observed droplet, the surface was measured. Each experiment was performed in triplicate.

### ThS assay

In a 384 well microplate with clear bottom (Greiner), samples were prepared in a total volume of 20 µL supplemented with 15µM Thioflavin S (ThS). The red shift of its emission spectrum upon binding with beta-sheets, was recorded quantitatively in a BioTek SynergyTM Mx plate reader by measuring emission at 490 nm after excitation at 450 nm. Sample was additionally visualized by a Leica DFC7000 GT camera connected to a Leica DMi8 microscope. Similarly, The ThS red shift of emission spectrum was visualized by a FITC filter.

### FRAP analysis of dynamics of LLPS droplets

Fluorescence recovery after photobleaching (FRAP) was used for assessing internal dynamics G3BP1-DPR condensates formed by LLPS. To this end, samples were prepared as described in the earlier paragraph. Next, pre-bleach images were recorded for 10 frames before the bleaching of droplets was performed by a laser shot for 50 ms. Recovery of the fluorescence was then captured for 200 frames. Each experiment was performed in triplicate.

### Dynamic light scattering (DLS)

To measure droplet size, dynamic light scattering (DLS) was performed on a DynaPro NanoStar (Wyatt). Before measuring, 40 µL of phase separating sample was administered to a disposable cuvette (Wyatt), surrounded by a buffer in a separate chamber. Intensity of scatter was then measured at a scattering angle of 95° at 25°C for 100 min, collecting 10 acquisitions of 8 sec. The analysis of the data was carried out by software package DYNAMICS 7.1.9. and hydrodynamic radius (Rh) was calculated. Each experiment was performed in duplicate.

### Measuring Kd of interactions by microscale thermophoresis (MST) and bio-layer interferometry (BLI)

For determining the Kds of G3BP1-RNA and G3BP-DPR interactions, two techniques, microscale thermophoresis (MST), and bio-layer interferometry (BLI), were used, as follows

#### Microscale thermophoresis

Microscale thermophoresis (MST) was carried out on a NanoTemper MonolithTM NT.115 instrument. Fluorescently labeled G3BP1 (see Methods) at 7 nM was titrated with different concentrations of polyU (Sigma Aldrich) in 50 mM HEPES, 100 mM NaCl, 0.05% Tween 20, pH 7.5, in a final volume of 20 µL in PCR tubes. Samples were loaded into MonolithTM NT.115 premium coated MST capillaries. The measurements were carried out at 50% MST power and 50% LED power. To fit data and calculate dissociation constants, the MO.Affinity Analysis software was used. Data were plotted with the GraphPad Prism software.

#### Bio-layer interferometry

Octet bio-layer interferometry (BLI) experiments were carried out in a buffer of 50 mM HEPES, 100 mM NaCl, pH 7.5, 0.05 % Tween 20, pH 7.5, on an Octet RED96 instrument. PolyU was loaded onto Streptavidin (ForteBio) sensors, followed by recording the baseline in buffer, then monitoring the association and dissociation of the protein. G3BP1 was used at concentrations 1-75 µM at 25 oC, at a shaking speed of 1000 rpm.

The BLI response signal was monitored first in a buffer for 120 sec, followed by loading for 300 sec, then washing for 120 sec. The baseline recorded for 120 sec was followed by association and dissociation for 600 and 900 sec, respectively. The dissociation constant (Kd) was estimated by a fitting response (nm) as Octet data Analysis Software 9.0. The final graph was generated by GraphPad Prism.

### Data collection for bioinformatics analysis

Interactors of R-DPRs were collected from various high-throughput screens (HTS). In PR100-FLAG immunoprecipitation, 177 interactors were found [45]. In PR50 and GR50 GFP immunoprecipitation, 194 high-confidence interactors were identified, and PR50-interactors and GR50-interactors were distinguished [26]. Finally, in a proximity-dependent biotin identification (BioID), interactors of PR100 (73), GR100 (83), and GA100 (82), were identified [46]. In the bioinformatic analysis, we have considered (i) all high-confidence R-DPR (PR + GR) interactors merged from the above three sources, (ii) PR and GR interactors separately, and (iii) GA interactors obtained from Liu et al. as negative controls (Suppl. Table S4) [46].

Of DPR interactors, we have also analyzed proteins identified in a HTS RNAi screen of genetic modifiers of GFP-GR50 toxicity in Drosophila melanogaster, i.e., enhancers and suppressors [26]. Human orthologs of 21 Drosophila melanogaster enhancers (with <25% viability) and 66 suppressors (with >48% viability) were obtained from Figure 2E of Lee et al. [26] (Suppl. Table S4).

The G3BP1 interactome along the line suggested in [35] was taken from the BioGRID database by only accepting interacting partners supported by low throughput or at least two high throughput evidence. A total of 135 proteins were obtained and analyzed (Suppl. Table S4).

The interactors of NPM1 were taken from the BioGRID database by only accepting interacting partners supported by low throughput or at least two high throughput evidence. The obtained list was complemented by the 132 NPM1 interactors identified in a pull-down assay performed as in [29], to finally arrive at 433 unique NPM1 interactors (Suppl. Table S4).

The stress granule (SG) proteome was obtained from RNAGranuleDB [55] by taking only Tier 1 SG proteins. The nucleolar proteome was derived from [56], by only accepting proteins with “Enhanced” or “Supported” nucleolar localization to gain a high-confidence dataset.

The human proteome (UP000005640) was obtained from UniProt [57] on 24.01.2023 totaling 20594 proteins

### Bioinformatics analyses

Proteins collected in the previous section were assessed for the presence/value of various features (Suppl. Table S5). LLPS propensity was predicted using DeePhase [47] for all proteins in the human proteome. A strict threshold of >0.75 was applied on the DeePhase score to obtain predicted LLPS drivers. The PFAM identifiers of 791 RNA-binding domains were obtained from EuRBPDB [58] and all proteins were tested for the presence of at least one of those RNA-binding domains (RBDs) according to their UniProt annotations. Also, all proteins were tested for the presence of at least two R-motifs (RxR or RxxR as described in [29]) separated by a maximum of 10 residues, and the presence of at least one di-RGG or tri-RG motif (RGG(X0-4)RGG and RG(X0-4)RG(X0-4)RG as defined previously [59]). Furthermore, all proteins were assessed for the presence of charged patches, i.e., sequence windows of 30 residues with a net charge ≤-10 (w30-10), or ≥+10 (w30+10). 145 proteins were excluded from the proteome analysis due to being shorter than 30 amino acids. The simultaneous presence of oppositely charged patches, and oppositely charged patches plus an RBD were also assessed for the proteins.These features were assessed for the different protein datasets explained in the previous section and the statistical significance of the fraction of proteins positive for a given feature was evaluated by comparing it to the respective proteome value using chi2 tests (Suppl. Table S6). We made multiple comparisons, eight features were compared between the different groups, therefore Bonferroni correction was applied on the significance levels: normally applied significance levels were reduced by tenfold for simplicity.

### Ultracoarse-grained multi-chain simulations

Ultracoarse-grained multi-chain Monte Carlo simulations were performed using the LASSI lattice-based simulation engine [60], similarly to earlier studies [34, 60, 61]. The monomeric G3BP1 structure was coarse-grained as 1 bead for the NTF2 domain, 2 beads for IDR1(-), 2 beads for IDR2, 1 bead for the RRM and 2 more beads for IDR3(+). The dimeric G3BP1 was coarse-grained as two protomers linked via their NTF2 beads. Both DPRs and RNA were modeled by 2-2 beads. All bead-to-bead distance was set to 1. Interactions potentials (measured in kT) were parametrized as in Suppl. Table S3. Bead-bead interactions extend only between neighboring lattice units. Monomer, dimer, trimer simulations were run in a cubic box of 10×10×10, while LLPS simulations were run in a cubic box of 50×50×50. A total number of 20 million Monte Carlo steps were performed for all simulations. Coarse-grain polymer architecture of the wild-type macromolecules (G3BP1 monomer, G3BP1 dimer, RNA and DPR) were set as outlined in Suppl. Figures S8 and S9.

## Abbreviations

ALS: amyotrophic lateral sclerosis
BLI: bio-layer interferometry
ΔIDR3-G3BP1: G3BP1 construct from which IDR3 is deleted
C9-ALS/FTD: ALS/FTD caused by C9orf72 repeat expansion
DLS: dynamic light scattering
DM2: myotonic dystrophy type 2
DPR: dipeptide repeat
EMSA: electrophoretic mobility shift assay
FITC: fluorescein isothiocyanate
FL: full-length
FRAP: fluorescence recovery after photobleaching
FTD: frontotemporal dementia
FUS: fused in sarcoma
G3BP1/2: Ras GTPase-activating protein-binding protein 1/2
GFP: green fluorescent protein
hnRNPA1: heterogeneous nuclear ribonucleoprotein A1
HTS: high-throughput screening
IDR: intrinsically disordered region
iMN: induced motor neuron
iMNs: induced motor neurons
IPTG: isopropylthio-β-galactoside
Kd: dissociation constant
LCD: low-complexity domain
LLPS: liquid-liquid phase separation
MST: microscale thermophoresis
NPM1: Nucleophosmin
NTF2: nuclear transport factor 2
NTF2L: NTF2-like (domain)
OD: optical density
PEG: polyethylene glycol
RAN-translation: repeat-associated non-AUG translation
RBP: RNA-binding protein
RBD: RNA-binding domain
Rh: hydrodynamic radius
RNAi: RNA interference
R-DPR: arginine-rich DPR
Rg: radius of gyration
Rh: hydrodynamic radius
RRM: RNA-recognition motif
SAXS: small-angle X-ray scattering
SCA36: spinocerebellar ataxia type 36
SG: stress granule
TDP-43: TAR DNA-binding protein 43
ThS: Thioflavin S
ThT: Thioflavin T
TIA1: cytotoxic granule-associated RNA-binding protein TIA1 (T-cell-restricted intracellular antigen-1)
UBAP2L: ubiquitin-associated protein 2-like
XDP: X-linked dystonia parkinsonism

## Supporting information

Table_S4_Datasets_used

## Acknowledgements

This work was supported by an EC H2020-WIDESPREAD-2020-5 Twinning grant (PhasAge, no. 952334) and EC H2020-MSCA-RISE Action grant (IDPfun, no. 778247, and grants K124670 and K131702 (to PT) and FK128133 and FK142285 (to RP) from the National Research, Development and Innovation Office (NKFIH), Hungary. The contribution of a VUB Strategic Research Program on Microfluidics (SRP51) at Vrije Universiteit Brussel (VUB, Brussels, Belgium, to MVN, DM and PT), an FWO PhD fellowship in strategic basic research (FWOSB77, to JA) is also acknowledged. TL is holder of a postdoctoral innovation mandate (grant no. HBC.2022.0194) by the Flanders Innovation & Entrepreneurship Agency (VLAIO). Furthermore, we personally thank Prof. Paul Taylor (St. Jude Children’s Research Hospital, Memphis, TN, USA) for providing the construct of full-length G3BP1.

## Supplementary material

**Figure S1:**
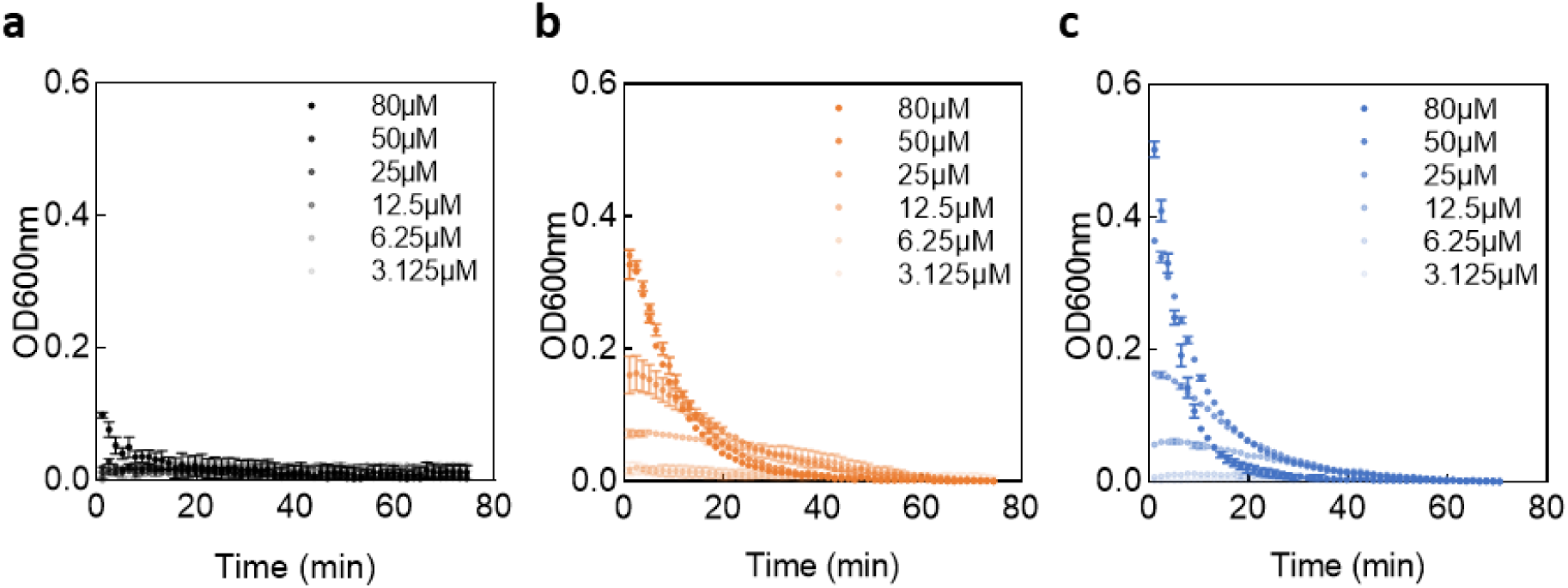
Turbidimetry of G3BP1 LLPS promoted by polyU, PR30 and GR30. OD at 600 nm (OD600nm) is recorded of polyU (a), PR30 (b) or GR30 (c), titrated with G3BP1 (going from 3.125 µM to 80 µM).

**Figure S2:**
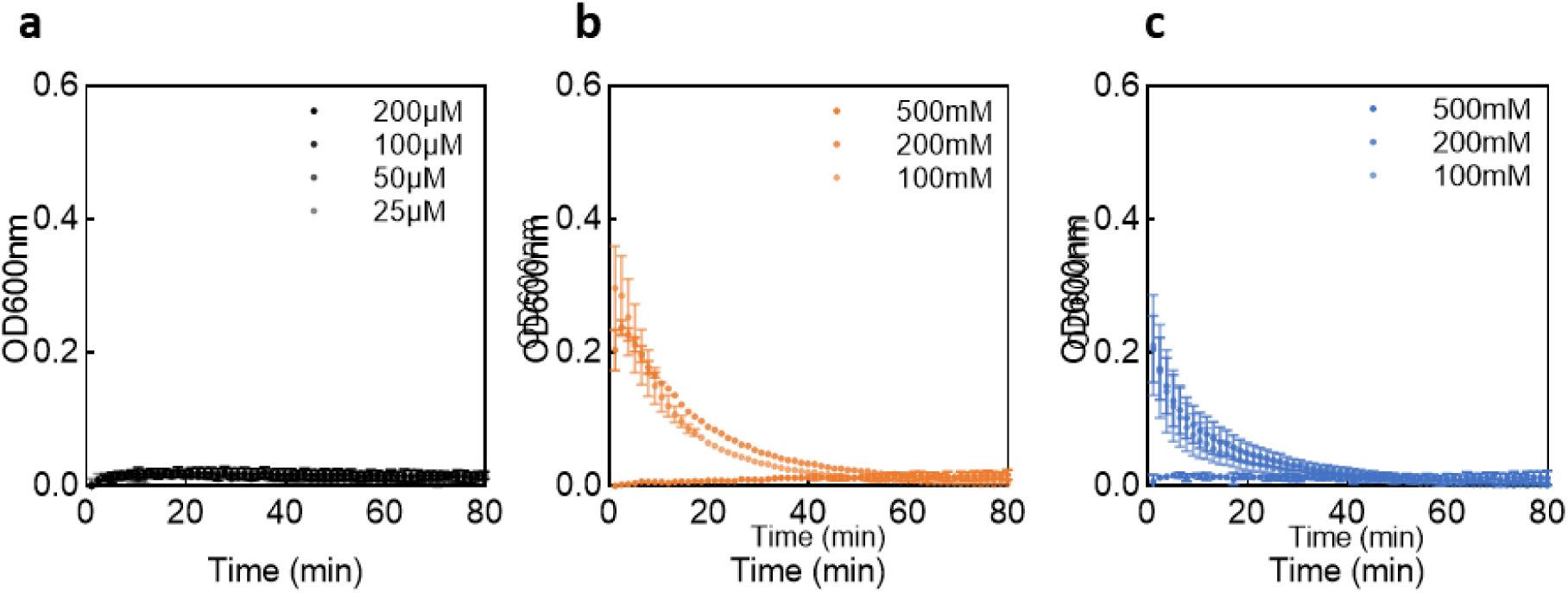
Turbidimetry of G3BP1 LLPS promoted by polyU, PR30 and GR30. OD at 600 nm (OD600nm) of G3BP1 recorded, upon titration with polyU (a), PR30 (b) or GR30 (c), concentrations going from 25 µM to 200 µM (polyU) or from 5 µM to 400 µM (PR30 and GR30).

**Figure S3:**
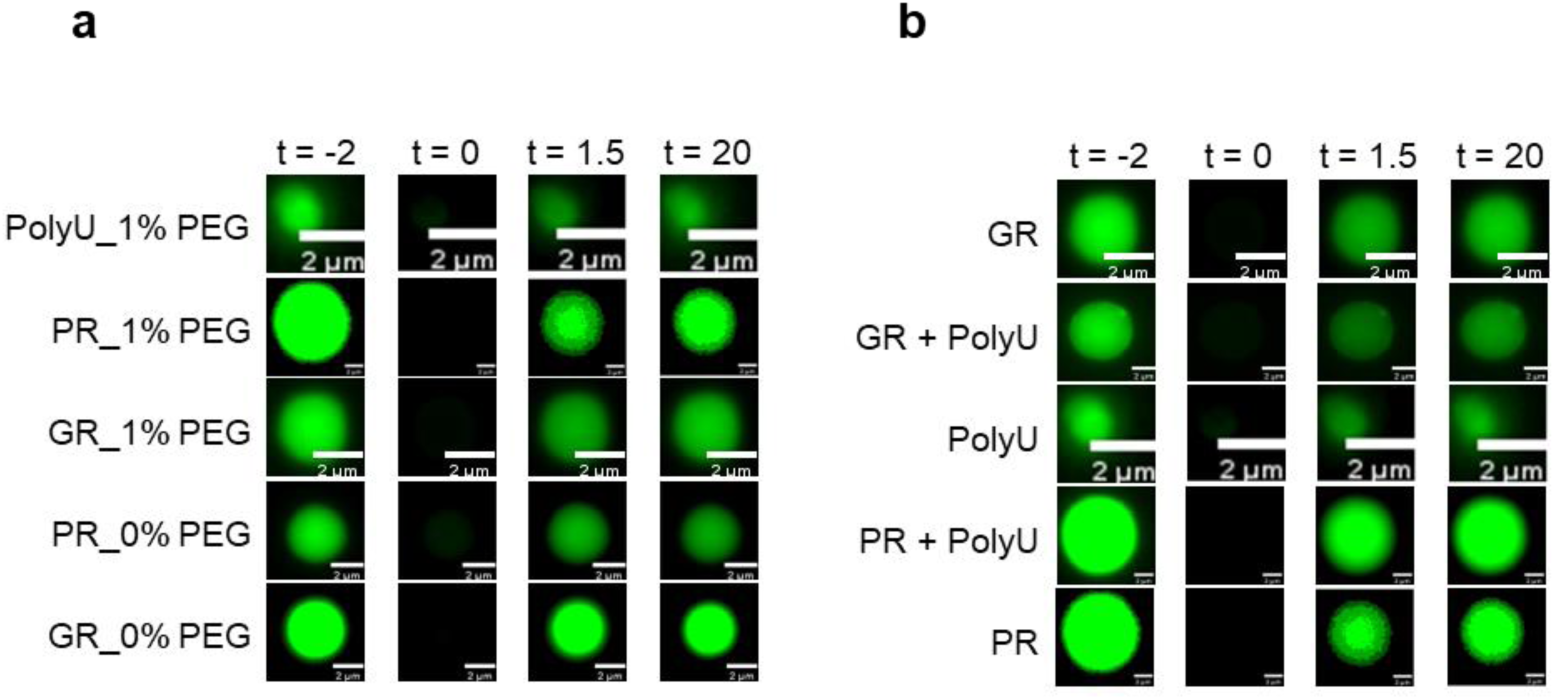
Microscopic images recorded during FRAP experiments. Pre-bleach (t = −2s), bleach (t = 0s), early post-bleach (t = 1.5s) and late post-bleach (t = 20s) images are reported. (a) G3BP1 droplets in the presence of polyU, GR30 and PR 30 in presence (top 3) or absence (bottom 2) of 1% PEG crowding agent. (b) G3BP1 droplets in the presence of GR30, GR30 plus polyU, polyU, PR30 plus polyU and PR30.

**Figure S4:**
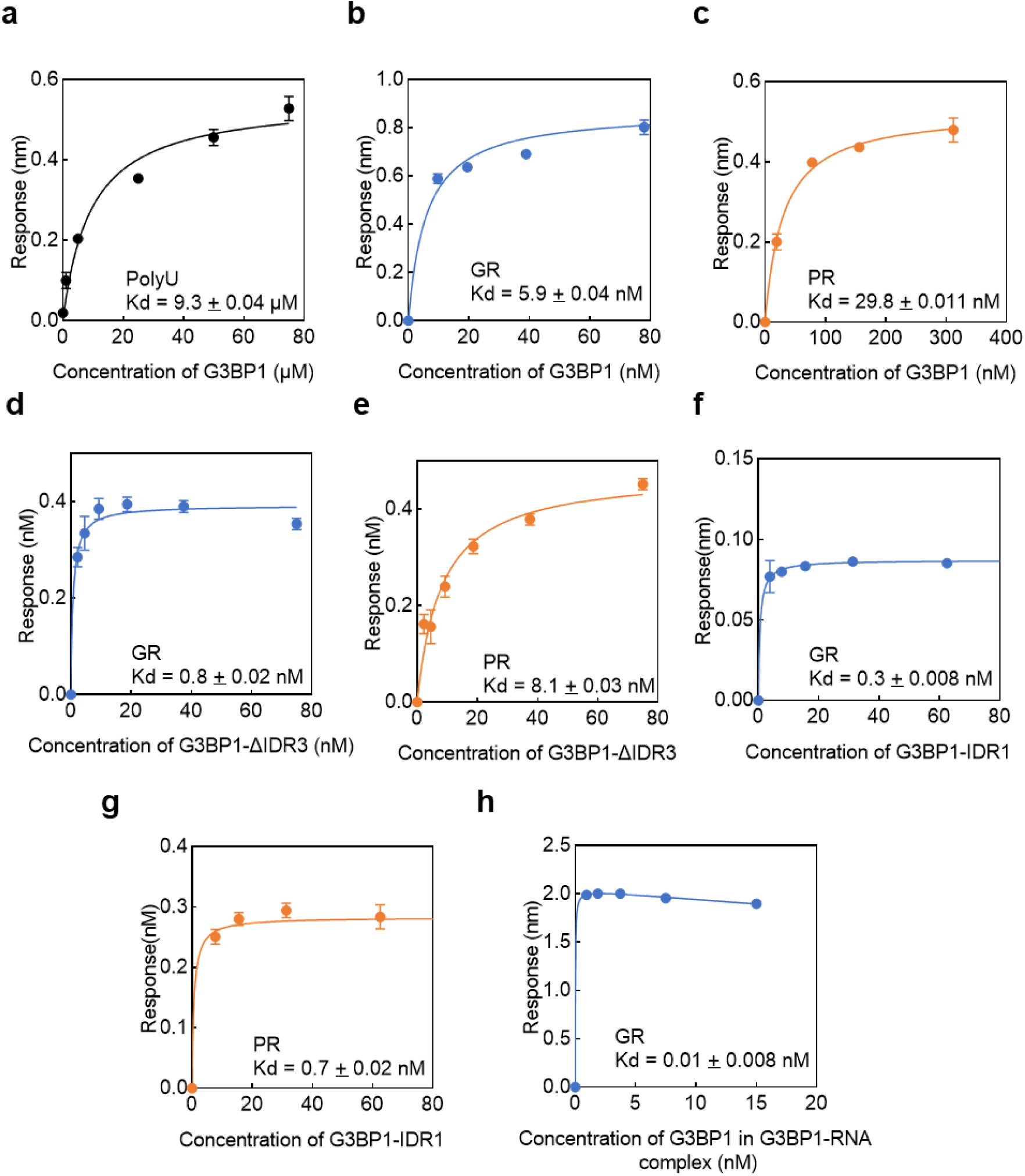
Determining Kd values of G3BP1 binding with biolayer interferometry (BLI) BLI response signal of G3BP1 (a, b, c), G3BP1-DIDR3 (d, e), or G3BP1-IR1 (f,g), added to the binding partner polyU (a), GR30 (b, d, f), or PR30 (c, e, g). BLI response signal of equimolar mixture of G3BP1-polyU added to the binding partner GR30 (h). The binding partner in all experiments is immobilized and titrated with the indicated G3BP1 construct. Kds were determined by fitting the data with a 1:1 binding model (Kds indicated on the panels and are also collected in Suppl. Table S2).

**Figure S5:**
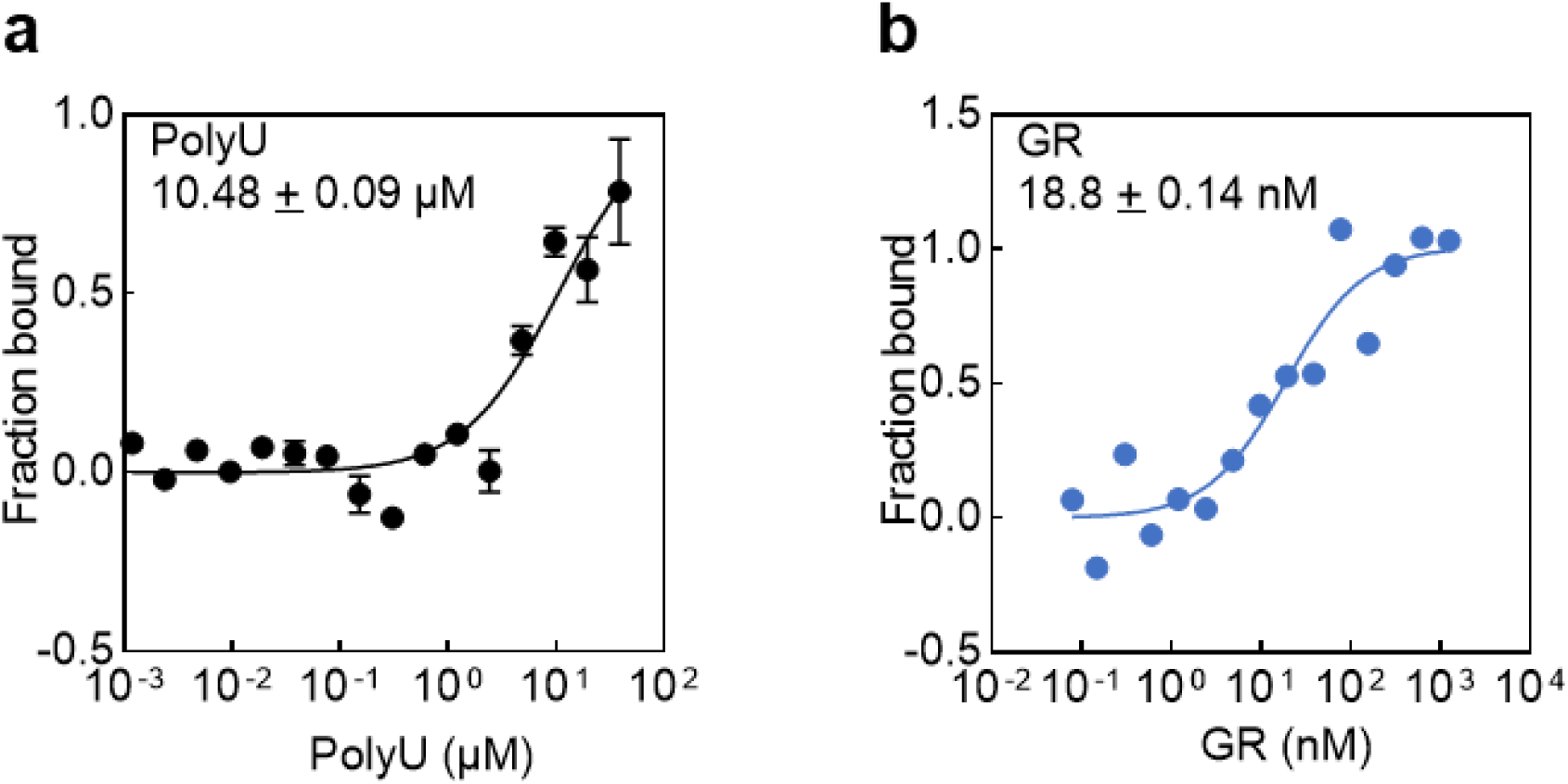
Determining Kd values of G3BP1 with microscale thermophoresis (MST) Cy5-labeled G3BP1was titrated with polyU (a) or GR30 (b), and the Kd of binding was determined by fitting the data with a 1:1 binding model. (Kds are indicated on the panels and are also collected in Suppl. Table S2).

**Figure S6:**
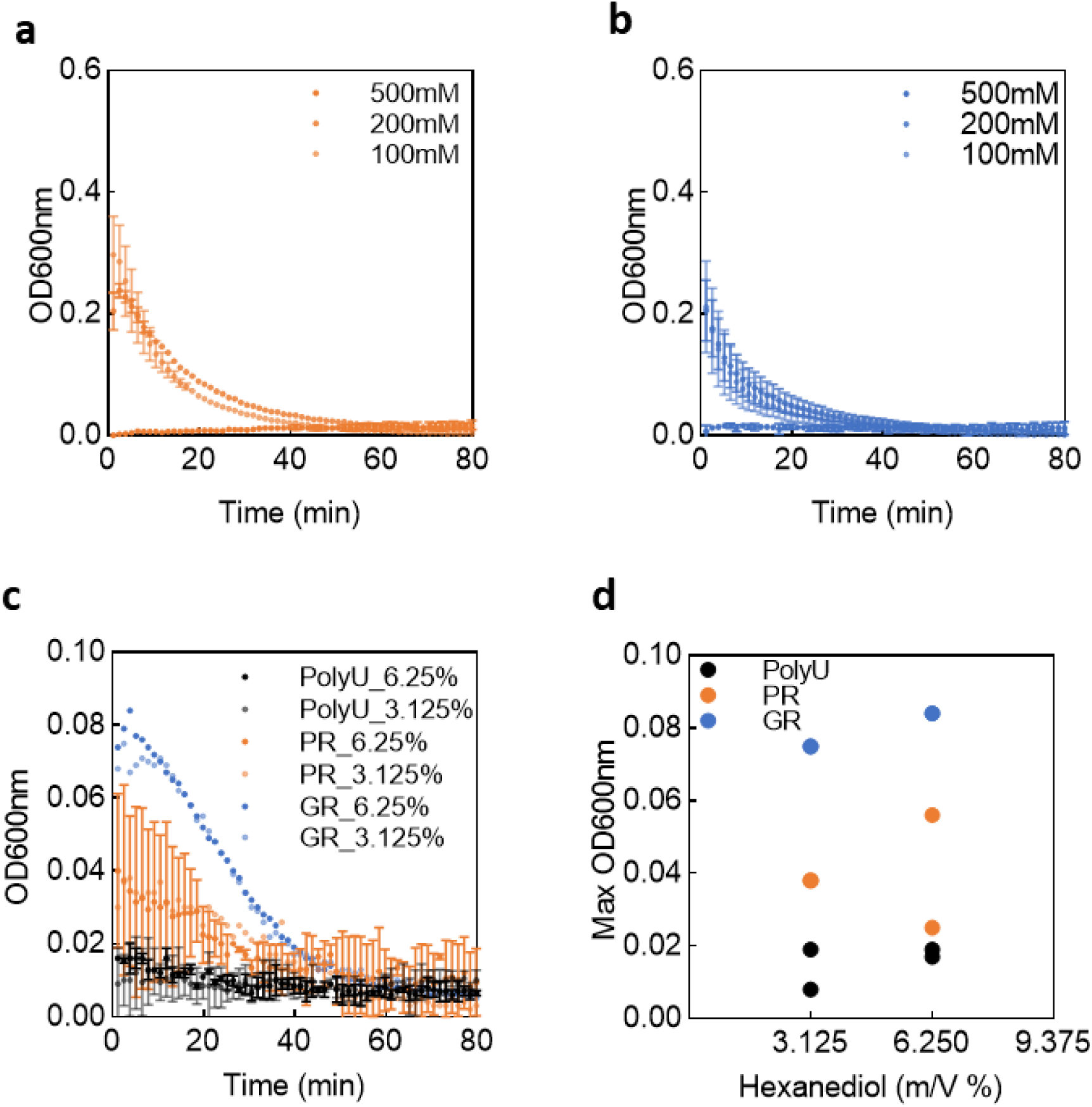
Effect of salt and 1,6-hexanediol on R-DPR - G3BP1 LLPS. Turbidity curves (OD600nm) of G3BP1 (25 µM), and PR30 (50 µM, a) or GR30 (50 µM, b) LLPS were recorded in the presence of increasing concentrations of salt (indicated). In similar experiments with GR30 (c), 1,6-hexanediol at increasing concentrations was added, and the maximum of the OD600nm curves for both PR30 and GR30 was also plotted as a function of 1,6-hexanediol concentration (d).

**Figure S7:**
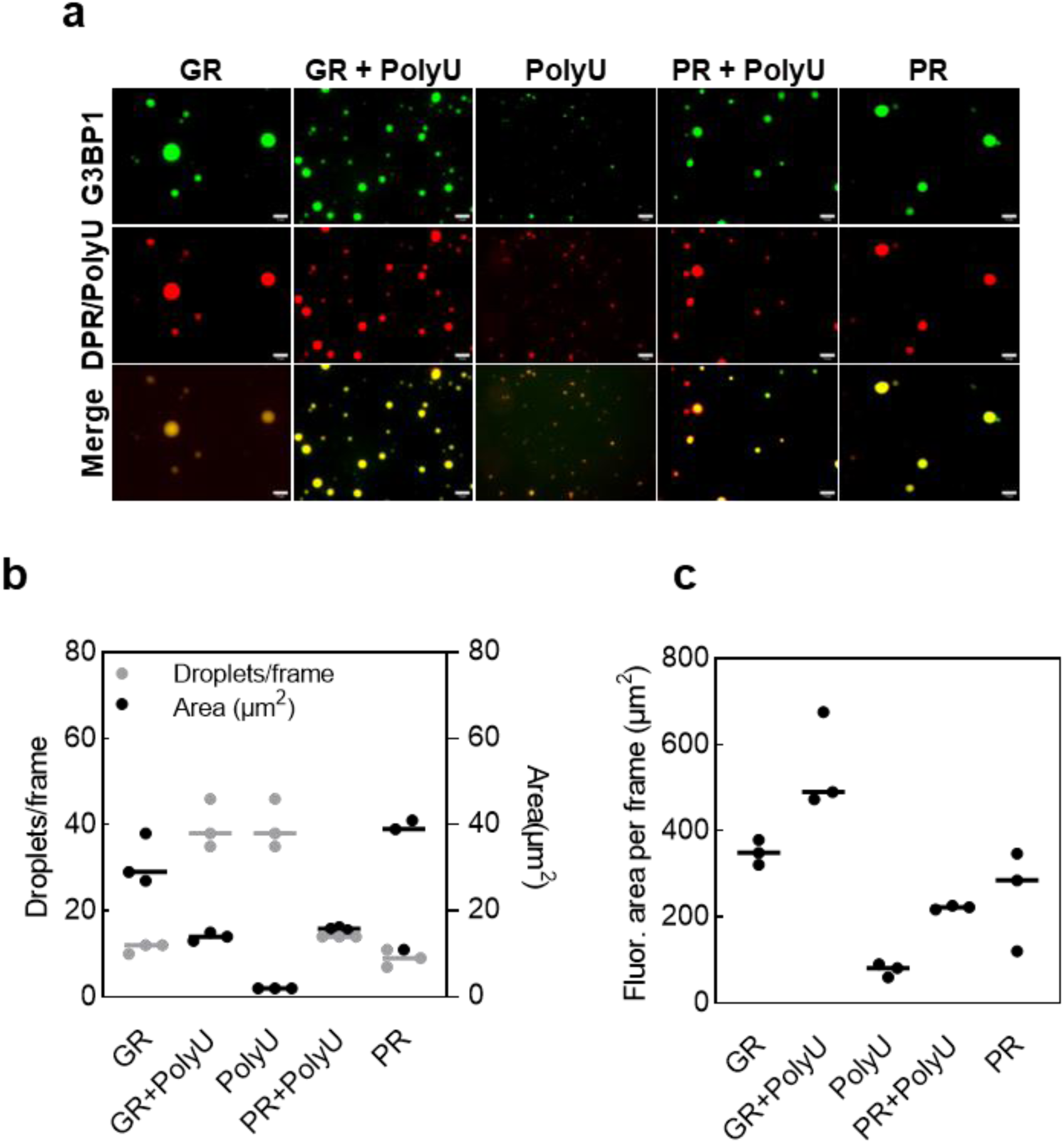
Cooperative effect of polyU and R-DPRs on G3BP1 LLPS. (a) Microscopic images of LLPS droplets of G3BP1 (60 µM) with polyU (50 µM), PR30 (50 µM) or GR30 (50 µM) alone, or polyU plus GR30, or polyU plus PR30 in combination. The scale bar on all images represents 10 µm. (b) G3BP1 droplet statistics of fluorescent images (panel a, repeated 3 times). Mean ± SD of the number of droplets within 3 observation areas and mean ± SD of the surface area of the droplets. (c) Total fluorescent surface area per frame (number of droplets x their average surface) on panel b.

**Figure S8:**
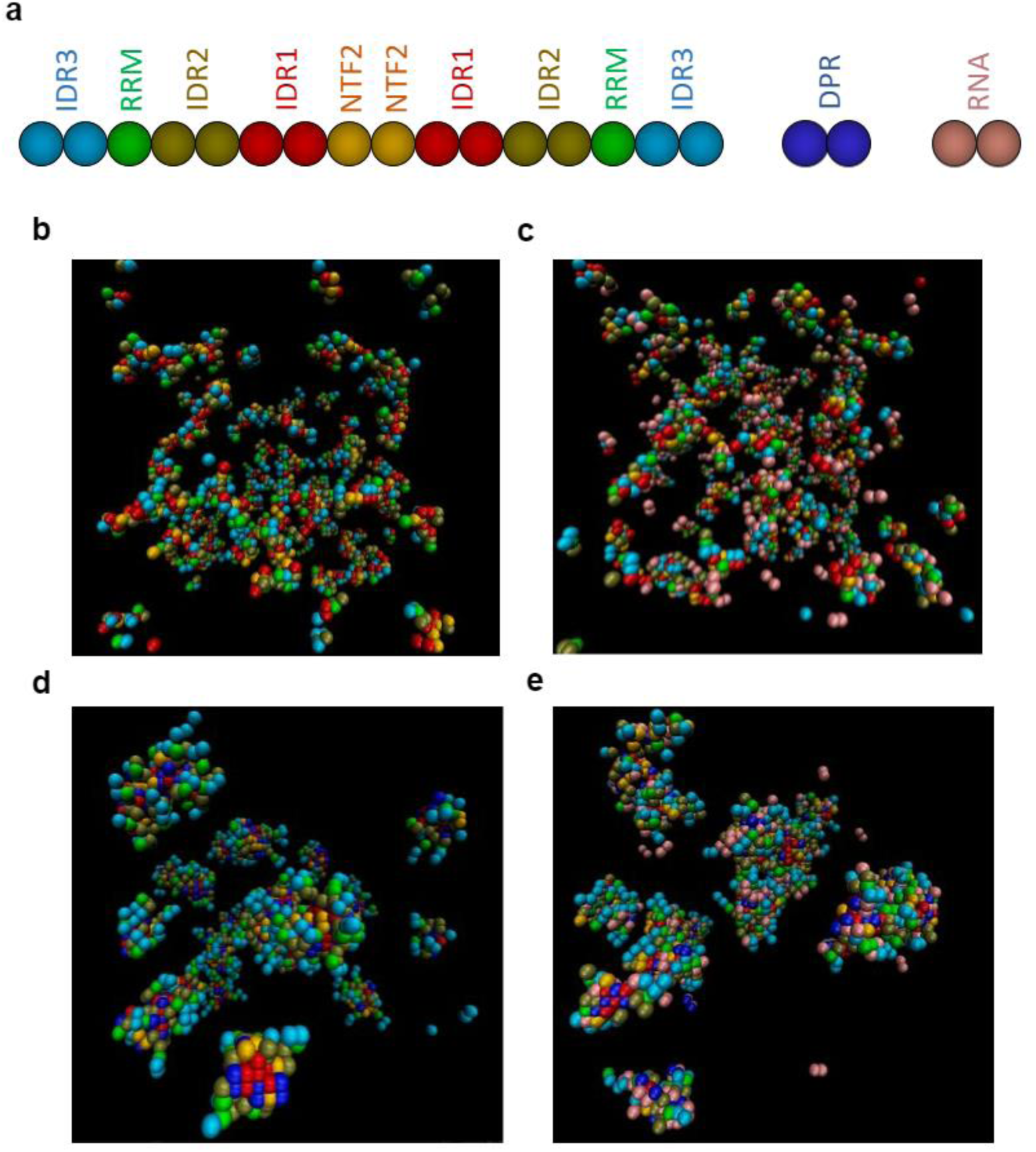
LLPS simulation of G3BP1 dimers and its partners from ultracoarse-grained multi-chain Monte Carlo simulations. 200 G3BP1 dimers (coarse-grained polypeptide architectures is shown, with colored beads or bead pairs matching domains, panel a) were simulated in the absence of partners (b), or in the presence of 200 RNA molecules (c), 200 DPR molecules (d), or 200 RNA and 200 DPR molecules (e).

**Figure S9:**
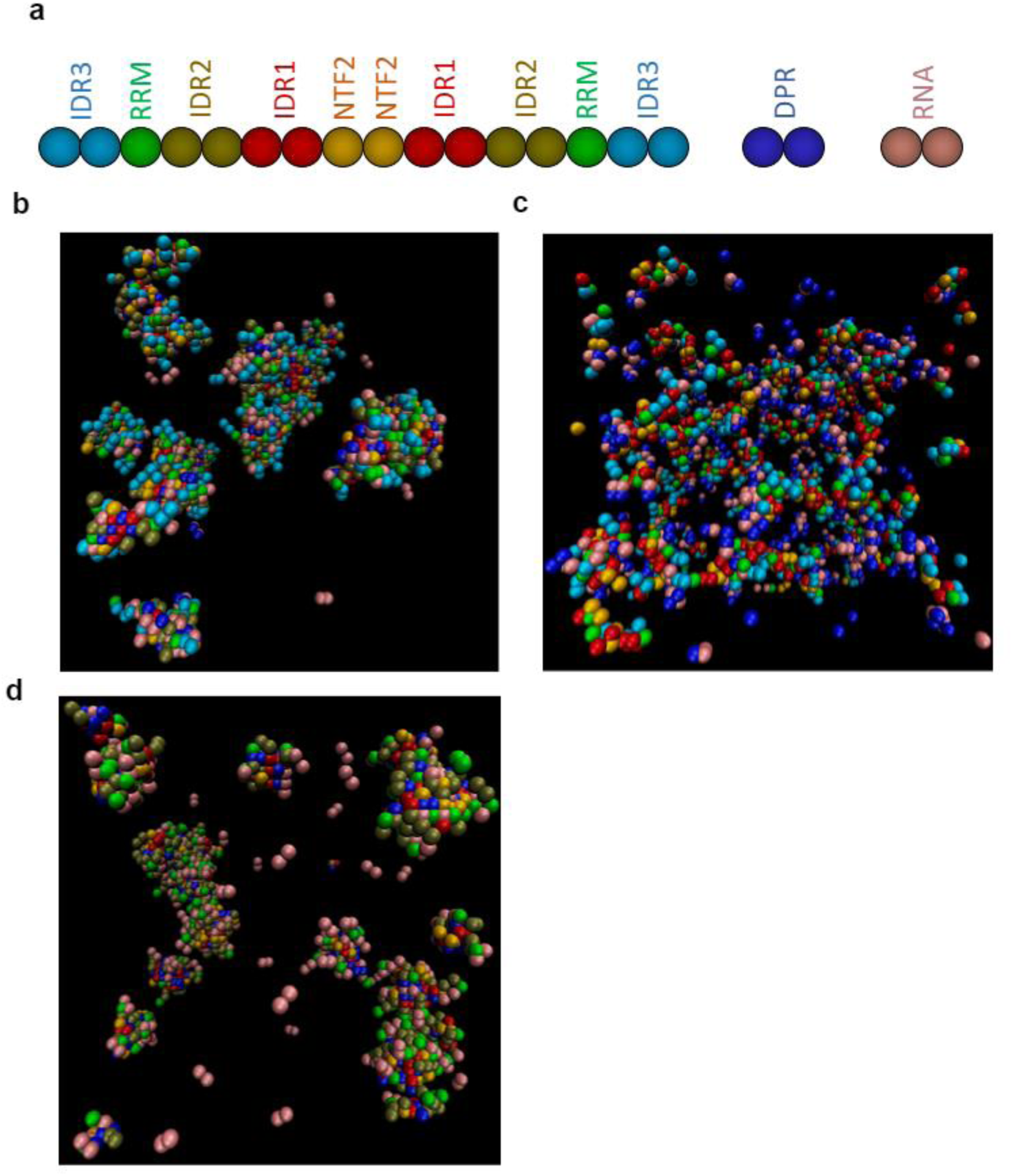
LLPS simulation of wild-type G3BP1 dimers and deletion constructs from ultracoarse-grained multi-chain Monte Carlo simulations. Ultracoarse-grained Monte-Carlo simulation (coarse-grained polypeptide architectures is shown, with colored beads or bead pairs matching domains, panel a) was carried out on 200 molecules of wt G3BP1 (b), G3BP1-ΔIDR1 (c) or G3BP1-ΔIDR3 (d), in the presence of 200 DPR and RNA molecules.

**Figure S10:**
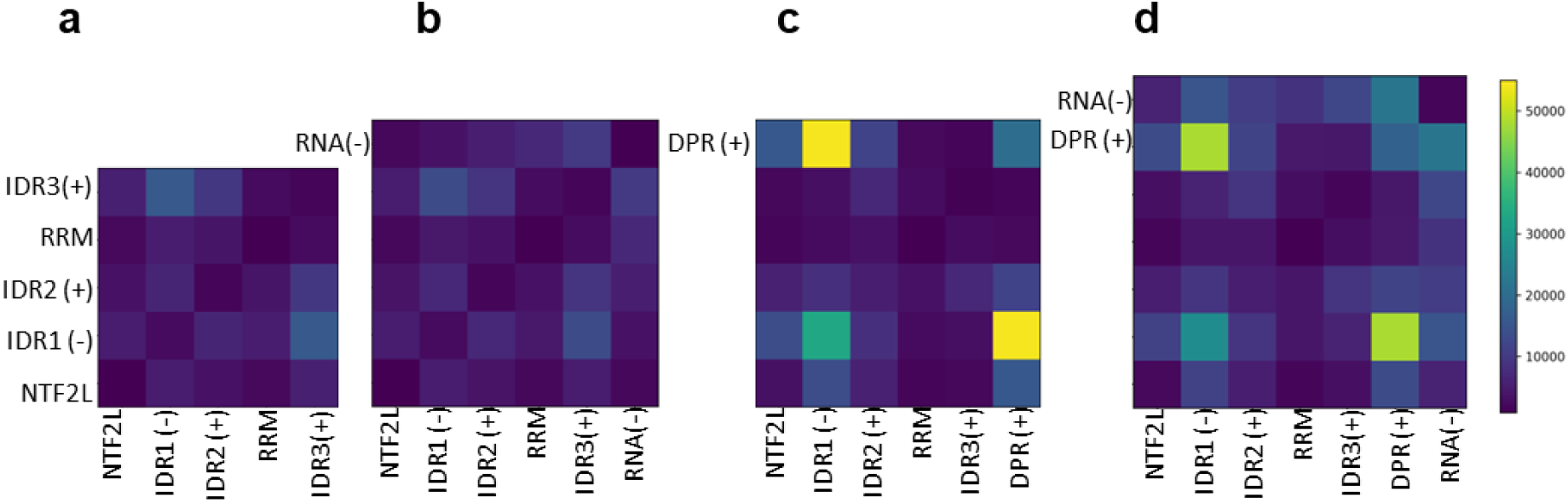
Region-region contact maps of G3BP1 and its partners from ultracoarse-grained multi-chain Monte Carlo simulations. Region-region contact map of G3BP1 simulated in the monomeric state (a), in complex with RNA (b), in complex with DPR (c), and in complex with both RNA and DPR (d).

**Figure S11:**
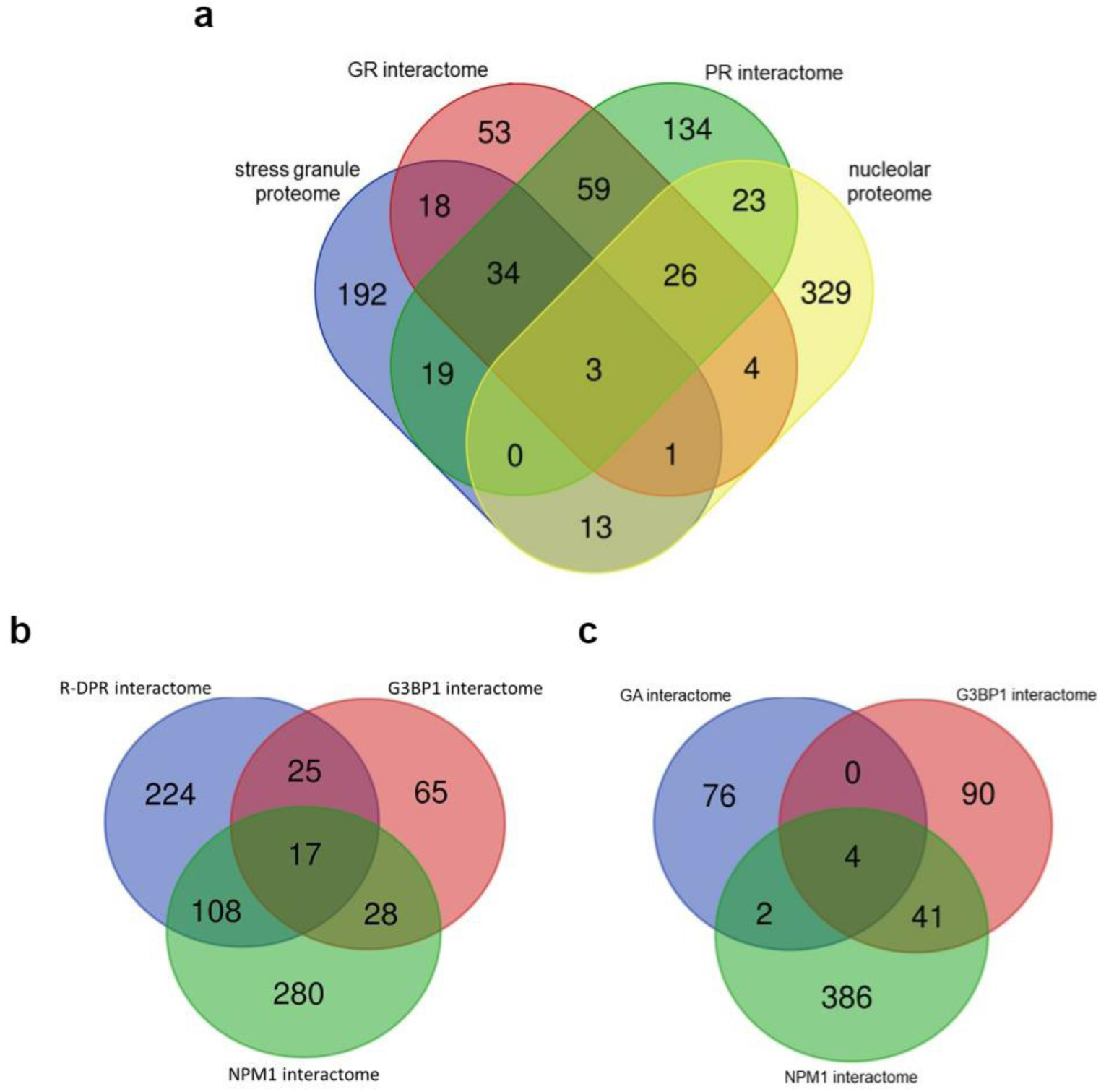
Comparison of the cellular interaction preferences of DPRs. Venn diagrams of the cellular interactomes of R-DPRs (poly-PR and poly-GR) and poly-GA: GR-and PR-interactomes are projected on the proteomes of stress granules and nucleoli (a), R-DPR interactome projected on the interactomes of NPM1 and G3BP1 (b), and poly-GA interactome projected on the interactomes of NPM1 and G3BP1 (c). (Data collected as described in Materials and methods).

**Table S1:** Supplementary data to FRAP experiments Half time and plateau with 95% confidence intervals of non-linear fit (GraphPad Prism software)

| **Sample** | **PEG** | **$t_{1/2}$** | **95% Confidence interval** | **Plateau** | **95% Confidence interval** |
| --- | --- | --- | --- | --- | --- |
| G3BP1 + PolyU | 1% | 0.29 | [0.20 ; 0.40] | 0.54 | [0.53 ; 0.55] |
| G3BP1 + GR | 1% | 1.07 | [1.00 ; 1.14] | 0.67 | [0.66 ; 0.67] |
| G3BP1 + PR | 1% | 0.22 | [0.17 ; 0.28] | 0.48 | [0.48 ; 0.49] |
| G3BP1 + GR | 0% | 0.56 | [0.44 ; 0.73] | 0.79 | [0.78 ; 0.80] |
| G3BP1 + PR | 0% | 0.34 | [0.29 ; 0.40] | 0.73 | [0.73 ; 0.74] |

**Table S2:** Binding data for G3BP1 variants and NPM1. Kd of binding of various partners, PR30, GR30, PR23 or S6C to G3BP1 variants or NPM1, as measured in this paper (with biolayer interferometry (BLI) or microscale thermophoresis (MST)) - or taken from the literature (measured by fluorescence anisotropy).

| Partner protein | Partner RNA | Kd (BLI) | Kd (MST) | Kd (Fluor. Anisotropy) | reference |
| --- | --- | --- | --- | --- | --- |
| G3BP1-FL |  |  |  |  |  |
| PR30 |  | 29.8 ± 0.011 nM |  |  | this paper |
| GR30 |  | 5.9 ± 0.044 nM | 18.8 ± 0.14 nM |  | this paper |
|  | PolyU | 9.3 ± 0.040 µM | 10.48 ± 0.092 µM |  | this paper |
|  | A60 | 2.0 µM |  |  | [34] |
| G3BP1-ΔIDR3 |  |  |  |  |  |
| PR30 |  | 8.1 ± 0.030 nM |  |  | this paper |
| GR30 |  | 0.8 ± 0.020 nM |  |  | this paper |
|  | A60 | 3.8 µM |  |  | [34] |
| G3BP1 – IDR1 |  |  |  |  |  |
| PR30 |  | 0.7 ± 0.016 nM |  |  | this paper |
| GR30 |  | 0.57 ± 0.0013 nM |  |  | this paper |
| G3BP1-PolyU Complex |  |  |  |  |  |
| GR30 |  | 0.01 ± 0.0081 nM |  |  | this paper |
| NPM1 |  |  |  |  |  |
| PR23 |  |  |  | 50.4 ± 7.5 nM | [30] |
| S6C |  |  |  | 15.1 ± 1.7 µM | [30] |

**Table S3:** Bead-bead interaction energies of the coarse-grained model.

| **Bead 1** | **Bead 2** | **Strength (kT)** | **Comment** |
| --- | --- | --- | --- |
| NTF2 | anything | 0.0 | No attraction |
| IDR1 | IDR1 | 1.5 | Medium strong repulsion |
| IDR1 | IDR2 | 0.5 | Weak repulsion |
| IDR1 | RRM | 0.0 | No attraction |
| IDR1 | IDR3 | -1.0 | Moderate attraction |
| IDR1 | RNA | 1.0 | Moderate repulsion |
| IDR1 | DPR | -5.0 | Very strong attraction |
| IDR2 | IDR2 | 0.0 | No attraction |
| IDR2 | RRM | 0.0 | No attraction |
| IDR2 | IDR3 | 0.0 | No attraction |
| IDR2 | RNA | 0.5 | Weak repulsion |
| IDR2 | DPR | 0.0 | No attraction |
| RRM | RRM | 1.0 | Moderate repulsion |
| RRM | IDR3 | 0.5 | Weak repulsion |
| RRM | RNA | -2.0 | Strong attraction |
| RRM | DPR | 0.1 | Very weak repulsion |
| IDR3 | IDR3 | 1.5 | Medium strong repulsion |
| IDR3 | RNA | -1.0 | Moderate attraction |
| IDR3 | DPR | 2.0 | Strong repulsion |
| RNA | RNA | 5.0 | Very strong repulsion |
| RNA | DPR | -3.0 | Strong attraction |
| DPR | DPR | 5.0 | Very strong repulsion |

**Table S4:** All protein datasets used in bioinformatics analyses. R-DPR-, poly-GA-, G3BP1-and NPM1 interactomes selected for this study are in the Supplementary Excel sheet: Table_S4_Datasets_used.xlsx Selection of data is described in the Materials and methods section “Data collection for bioinformatics analysis”, and in the header of the table.

**Table S5:**
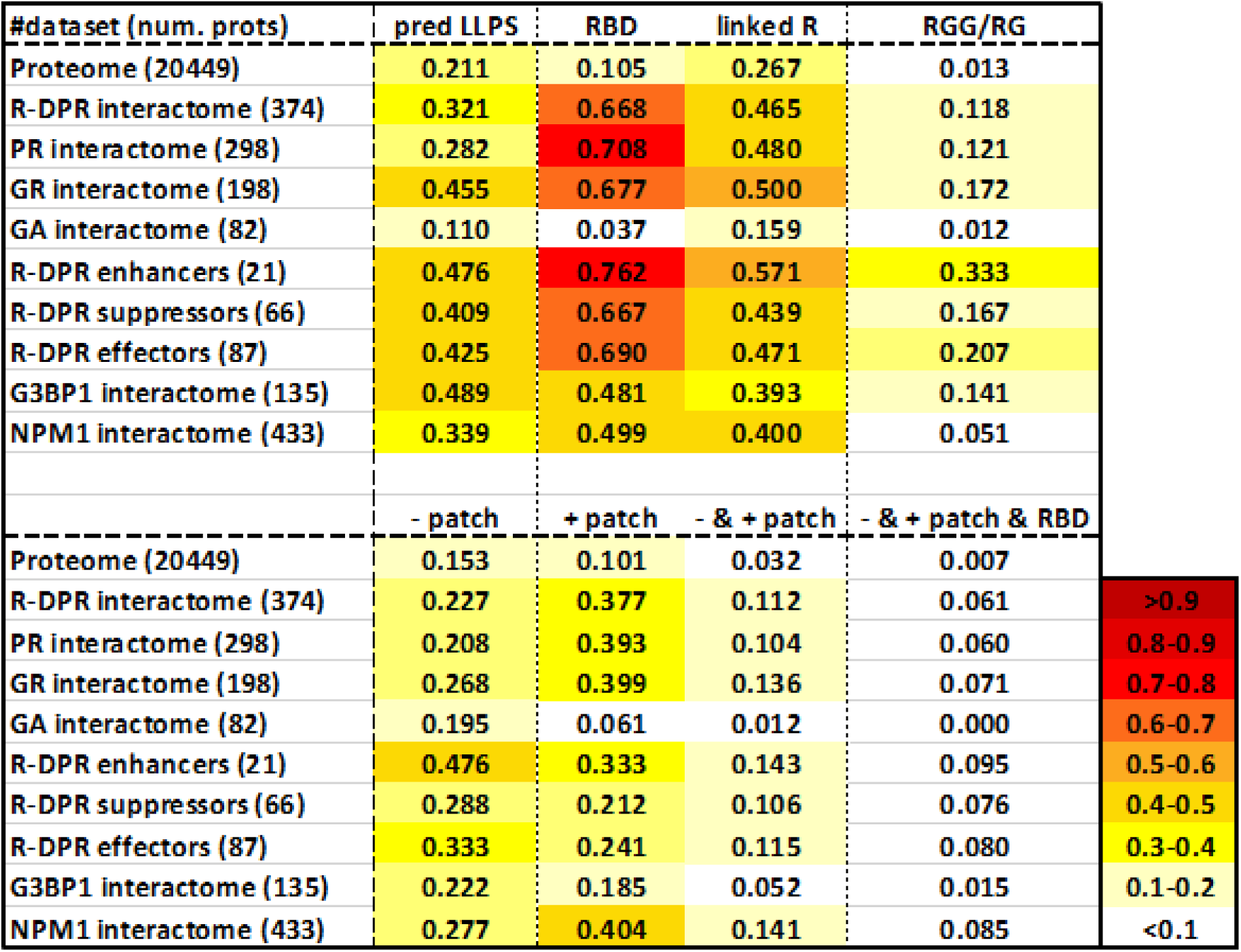
Fraction of proteins among the interaction partners of R-DPRs, G3BP1 and NPM1 for various features. The entire human proteome, and interactomes of R-DPRs (PR, GR and GA), proteins that enhance or suppress R-DPR toxicity (enhancer, suppressor, and their sum, “effectors”), and interactomes of G3BP1 and NPM1 were identified and analyzed for various features: LLPS propensity with the DeePhase predictor (at a threshold 0.75), the presence of an RNA-binding domain (RBD), two linked R motifs or an RGG/RG motif, a high local charge ("-patch": at least –10 net charge within a window of 30 residues, or “+ patch”: at least +10 net charge within a window of 30 residues), or a combination of – & + patch, or a – & + patch & RBD, within the same protein. Color coding indicates the fraction of proteins with the particular feature.

**Table S6:** Statistical evaluation of the differences between the human proteome and the interactomes of DPRs, NPM1 and G3BP1. The number of proteins positive for a given feature (columns) in a particular protein group (rows) was compared to the respective proteome value using chi2 tests. In each cell, the upper value is the calculated chi2 value, the middle value is the corresponding p-value, and the last row shows the significance levels. Bonferroni significance level correction was applied due to comparing multiple features between the groups: P< 0.005 *, p< 0.001 **, p< 0.0001 ***, n.s.: not significant.

| #dataset (num. prots) | pred. LLPS | RBD | linked R | RGG/RG | - patch | + patch | - & + patch | - & + patch & RBD |
| --- | --- | --- | --- | --- | --- | --- | --- | --- |
| Proteome (20449) | - | - | - | - | - | - | - | - |
| R-DPR interactors (374) | 27.7264;<br><0.00001;<br>*** | 1289.3901;<br><0.00001;<br>*** | 76.5873;<br><0.00001;<br>*** | 327.9434;<br><0.00001;<br>*** | 16.2349;<br>0.000056;<br>*** | 319.9643;<br><0.00001;<br>*** | 80.0267;<br><0.00001;<br>*** | 164.4141;<br><0.00001;<br>*** |
| PR interactors (298) | 9.1787;<br>0.002448;<br>* | 1172.3973;<br><0.00001;<br>*** | 70.0905;<br><0.00001;<br>*** | 276.2533;<br><0.00001;<br>*** | 7.0828;<br>0.007783;<br>n.s. | 283.5215;<br><0.00001;<br>*** | 51.1336;<br><0.00001;<br>*** | 125.3247;<br><0.00001;<br>*** |
| GR interactors (198) | 71.387;<br><0.00001;<br>*** | 696.8082;<br><0.00001;<br>*** | 55.517;<br><0.00001;<br>*** | 395.6777;<br><0.00001;<br>*** | 20.3051;<br><0.00001;<br>*** | 195.7298;<br><0.00001;<br>*** | 70.8762;<br><0.00001;<br>*** | 117.8794;<br><0.00001;<br>*** |
| GA interactors (82) | 5.0541;<br>0.024568;<br>n.s. | 4.0874;<br>0.043203;<br>n.s. | 4.9416;<br>0.026218;<br>n.s. | 0.0033;<br>0.954168;<br>n.s. | 1.1292;<br>0.287939;<br>n.s. | 1.4486;<br>0.228754;<br>n.s. | 1.0304;<br>0.310073;<br>n.s. | 0.3209;<br>0.571097;<br>n.s. |
| R-DPR enhancers (21) | 8.8981;<br>0.002855;<br>* | 96.6993;<br><0.00001;<br>*** | 9.9627;<br>0.001597;<br>* | 169.3654;<br><0.00001;<br>*** | 16.9504;<br>0.000038;<br>*** | 12.5114;<br>0.000404;<br>** | 8.4067;<br>0.003738;<br>* | 23.765;<br><0.00001;<br>*** |
| R-DPR suppressors (66) | 15.6457;<br>0.000076;<br>*** | 222.6859;<br><0.00001;<br>*** | 10.0678;<br>0.001509;<br>* | 122.8362;<br><0.00001;<br>*** | 9.3016;<br>0.002289;<br>* | 9.0204;<br>0.00267;<br>* | 11.8357;<br>0.000581;<br>** | 45.4679;<br><0.00001;<br>*** |
| R-DPR effectors (87) | 24.1558;<br><0.00001;<br>*** | 318.3821;<br><0.00001;<br>*** | 18.6478;<br>0.000016;<br>*** | 258.0031;<br><0.00001;<br>*** | 21.9377;<br><0.00001;<br>*** | 18.9914;<br>0.000013;<br>*** | 19.579;<br><0.00001;<br>*** | 68.4764;<br><0.00001;<br>*** |
| G3BP1 interactors (135) | 63.1598;<br><0.00001;<br>*** | 205.4008;<br><0.00001;<br>*** | 10.9712;<br>0.000925;<br>** | 174.2574;<br><0.00001;<br>*** | 5.0308;<br>0.024901;<br>n.s. | 10.63;<br>0.001113;<br>* | 1.7666;<br>0.183807;<br>n.s. | 1.2208;<br>0.269202;<br>n.s. |
| NPM1 interactors (433) | 44.0236;<br><0.00001;<br>*** | 731.6825;<br><0.00001;<br>*** | 39.7693;<br><0.00001;<br>*** | 49.8575;<br><0.00001;<br>*** | 52.6408;<br><0.00001;<br>*** | 448.2062;<br><0.00001;<br>*** | 170.6532;<br><0.00001;<br>*** | 395.3676;<br><0.00001;<br>*** |

**Table S7:** R-DPR interactors enriched with switch features. Of R-DPR interactors, the ones are enlisted which have a negative patch (≥-10 charge in a window of 30), a positive patch (≥+10/30), and an RBD within the same protein (cf. Fig. 4e). Of the 23 hits, the UniProt accession code (AC), gene name, UniProt disease link, membraneless organelle (MLO) localization (as indicated in PhaSepDB database) and LLPS propensity (as indicated in PhaSepDB database) are shown.

| UniProt AC | Gene name | Protein name | Disease link (UniProt) | MLO localization (PhaSepDB) | LLPS (PhaSepDB) |
| --- | --- | --- | --- | --- | --- |
| O75179 | **ANKRD17** | Ankyrin repeat domain-containing protein 17 | Chopra-Amiel-Gordon syndrome (CAGS) | SG, P-body | No |
| Q9NVP1 | **DDX18** | ATP-dependent RNA helicase DDX18 | - | Nucl., P-body | No |
| Q96GQ7 | **DDX27** | Probable ATP-dependent RNA | - | Nucl., P-body | No |
| Q9BQ39 | **DDX50** | ATP-dependent RNA helicase DDX50 | - | Nucl., SG, P-body | No |
| Q6P158 | **DHX57** | Putative ATP-dependent RNA helicase | - | Nucl., SG, spliceosome | No |
| O60841 | **EIF5B** | Eukaryotic translation initiation factor | - | Nucl., SG | No |
| Q8IY81 | **FTSJ3** | pre-rRNA 2'-O-ribose RNA methyltransferase FTSJ3 | - | Nucl., SG | No |
| Q9BVP2 | **GNL3** | Guanine nucleotide-binding protein- | - | Nucl. | No |
| Q9BZE4 | **GTPBP4** | GTP-binding protein 4 | - | Nucl., P-body | No |
| B7ZW38 | **HNRNPCL3** | Heterogeneous nuclear ribonucleoprotein C-like 3 | - | Not known | No |
| Q00839 | **HNRNPU** | Heterogeneous nuclear ribonucleoprotein U | Developmental and epileptic encephalopathy 54 (DEE54) | Nucl., Nuclear body, Nuclear speckle, P-body, Spindle apparatus, Spliceosome, SMN complex, Nuclear stress body, IMP1 ribonucleoprotein granule | No |
| Q6PKG0 | **LARP1** | La-related protein 1 | - | SG | No |
| Q9BXY0 | **MAK16** | Protein MAK16 homolog | - | Nucl., Sam68 nuclear body | No |
| O00566 | **MPHOSPH10** | U3 small nucleolar ribonucleoprotein protein MPP10 | - | Nucl. | No |
| P19338 | **NCL** | Nucleolin | - | Nucl., Cajal body, IMP1 ribonucleoprotein granule | YES |
| P46087 | **NOP2** | Probable 28S rRNA (cytosine(4447)-C(5))-methyltransferase | - | Nucl., P-body | No |
| Q5T8P6 | **RBM26** | RNA-binding protein 26 | - | Stress granule, Nuclear speckle | No |
| Q9GZR2 | **REXO4** | RNA exonuclease 4 | - | Nucl. | No |
| Q9Y3A4 | **RRP7A** | Ribosomal RNA-processing protein 7 homolog A | Microcephaly 28, primary, autosomal recessive (MCPH28) | Nucl. | No |
| Q96T21 | **SECISBP2** | Selenocysteine insertion sequence-binding protein 2 | Thyroid hormone metabolism, abnormal, 1 (THMA1) | SG | No |
| Q2NL82 | **TSR1** | Pre-rRNA-processing protein TSR1 | - | - | No |
| P26368 | **U2AF2** | Splicing factor U2AF 65 kDa subunit | - | Nucl., Nuclear body, Nuclear speckle, SG, Spliceosome, Paraspeckle | YES |
| Q9NQZ2 | **UTP3** | Something about silencing protein 10 | - | Nucl. | No |

